# Cortical substrates of perceptual confusion between pitch and timbre

**DOI:** 10.1101/2025.02.19.639197

**Authors:** Yongtian Ou, Emily J. Allen, Kendrick N. Kay, Andrew J. Oxenham

## Abstract

Pitch and timbre, two fundamental perceptual attributes of sound, are critical for both speech and music. Although governed by different acoustic parameters, pitch and brightness (one of the primary dimensions of timbre) can be perceptually confused. Here we identify potential neural substrates of this well-known illusion by independently manipulating pitch and brightness while measuring fMRI responses. First, we show that the preferred pitch of individual voxels within auditory cortex decreases systematically as stimulus brightness increases, and vice versa, consistent with predictions from perceptual confusion. Second, we demonstrate that pitch and brightness share a common tuning gradient across auditory cortex, implying a shared trajectory of cortical activation for changes in each dimension. The pattern of results at both local and global cortical levels are consistent with the principle of efficient coding, with the brain exploiting correlations between pitch and brightness in natural sounds via a shared neural code.

## INTRODUCTION

Pitch and timbre are commonly regarded as two distinct perceptual features of sound. In music, pitch defines melody and harmony, whereas timbre refers to the sound quality (e.g., of different musical instruments)^1^. In speech, pitch helps define prosody and, in some languages, semantic content, whereas timbre determines voice quality and is what allows us to distinguish between different speech sounds, such as the vowels “ah” and “oo”. Pitch and timbre also have different acoustic correlates. The pitch of a sound is determined primarily by its repetition rate, or fundamental frequency (F0)^2^. The timbre of a sound depends on various physical properties, including its spectral centroid (Fc), i.e., the center of mass in its energy distribution across frequencies, which is closely related to a key timbral attribute known as brightness^3,4^. Brightness is one of the most salient aspects of timbre, having been consistently identified as a primary factor in multidimensional scaling studies^5,6^.

Although they are treated as separate dimensions and are based on different acoustic properties^7^, pitch and brightness exhibit intriguing interdependencies in perception. For example, two tones are presented sequentially, a small increase in brightness from the first to the second can be mistaken for an increase in pitch^8^. Similarly, when pitch and brightness vary simultaneously, listeners’ estimates of the pitch interval are larger for a given change in F0 when the brightness varies in the same direction (congruent) than when it moves in the opposite direction (incongruent)^9^. Moreover, listeners’ judgments of the direction of pitch and brightness changes are faster and more accurate in congruent than in incongruent conditions^10–12^. Together, these findings demonstrate that changes in pitch and timbral brightness can be perceptually confused with each other.

To provide a mechanistic explanation for this phenomenon, one hypothesis posits that the perceptual confusion reflects a lack of complete separability between the two dimensions, so that changes in one affect the encoding of the other. Such interactions could lead to greater sensitivity when pitch and timbre vary in the same direction (as found in both speech^13^ and music^14^) than when they vary in opposite directions, in line with expectations based on efficient coding^15–18^. An alternative account is that pitch and brightness are encoded separately and independently but misattribution of changes in one feature for changes in the other leads to poorer performance when the dimensions vary in opposite directions. These two hypotheses are difficult to disentangle using only behavioral measures, but neural evidence may offer critical insights, as data can be acquired without requiring participants to make perceptual judgments.

Neural populations in the auditory cortex that are sensitive to pitch and brightness are largely co-localized, as demonstrated across multiple spatial scales of measurement. In single-unit recordings from ferrets, a substantial proportion of neurons in the auditory core and belt respond to both pitch and brightness, as well as sound source location^19,20^. Human fMRI studies similarly show that sensitivity to pitch and brightness variations are largely localized to the same cortical regions^21^. Evidence from human electrocorticography (ECoG) further reveals electrodes that encode both features jointly, as well as electrodes selective for only one^22,23^. These studies establish the spatial proximity of pitch-and brightness-sensitive neural populations, which is generally consistent with perceptual pitch-brightness confusion. However, it is not known whether changes in one dimension alter the neural tuning for the other dimension. Such changes would imply interactions in neural encoding, which could potentially increase the discriminability of congruent changes while decreasing the discriminability of incongruent changes, as observed in perceptual experiments.

This study sought to examine potential interactions between pitch and brightness in human auditory cortex to determine whether they are observed and, if so, whether they are consistent with the observed perceptual interference effects. By systematically measuring cortical responses to stimuli with fully crossed pitch and brightness levels, we were able to examine whether cortical tuning to one dimension is affected by the other. Our findings reveal robust interactions between representations of pitch and brightness within individual voxels in the same direction as the perceptual interference effects observed behaviorally. In addition, we observed overlapping and consistent preference gradients for both dimensions across auditory cortex in all of the brains studied that are also in a direction consistent with the perceptual effects. Taken together, our findings reveal a potential neural substrate for the long-recognized but poorly understood perceptual confusion between pitch and timbral brightness.

## RESULTS

### Perceptual confusion between pitch and timbral brightness

To confirm and quantify the perceptual confusion between pitch and brightness in our stimuli, we measured the effect of pitch changes (manipulated by varying the fundamental frequency, F0) on judgments of brightness changes (manipulated by varying the spectral centroid, Fc), and *vice versa*. Participants performed a pitch discrimination task and a brightness discrimination task, in which they were presented with two consecutive tones and asked to judge either which tone had a higher pitch (pitch discrimination) or which tone had a brighter timbre (brightness discrimination). First, the difference limens (DLs) for pitch and brightness alone were measured in each participant individually. Next, in the pitch discrimination task, the two tones differed by one F0 DL (DL_F0_) while also varying from-2 to +2 DL_Fc_, in steps of 1 DL_Fc_. In the brightness discrimination task, the two tones differed by one DL_Fc_ while also varying from-2 to +2 DL_F0_. Positive values indicate congruent changes, meaning that both the target and non-target dimension changed in the same direction between the two tones. If there were no confusion between pitch and brightness, changes in the non-target dimension should have no effect on performance. In contrast, we observed a significant positive relationship between discrimination performance and changes in the non-target dimension, consistent with directional perceptual confusion between the two dimensions (Fig. 1A).

**Figure 1.**
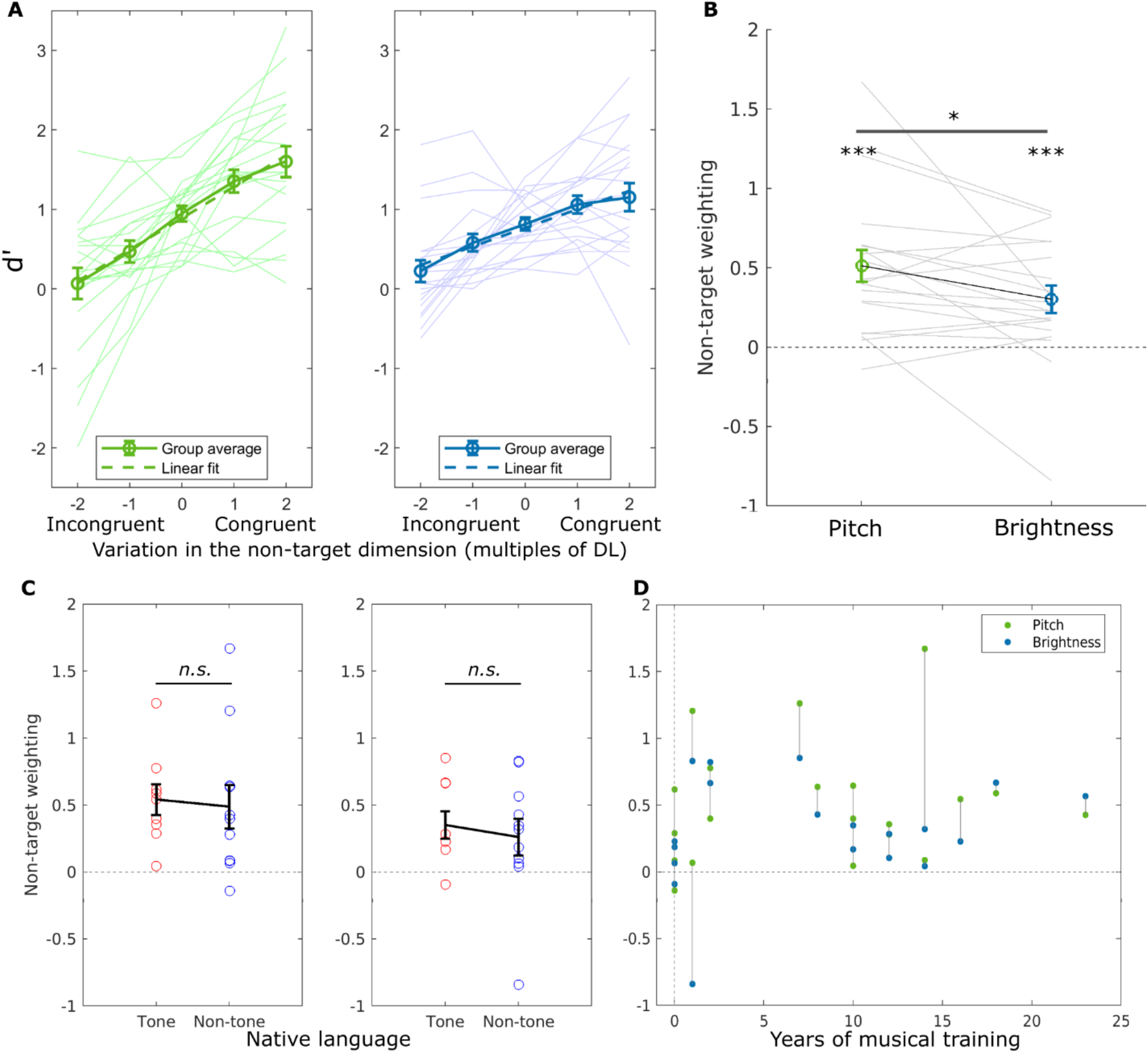
Behavioral confusion between pitch and timbral brightness. **(A)** Discrimination of the pitch (left) and brightness (right) as a function of the variation in the non-target dimension. A positive slope implies that increases in pitch or brightness are confused for increases in the other dimension. Thick solid line: group average; thick dashed line: linear fit of the group average; thin solid lines: individual data. **(B)** Perceptual weight of the non-target dimension on judgments of the target dimension, obtained from individual participants’ linear fits in (A). Black line connects group average perceptual weights for pitch and brightness discrimination. Grey lines connect perceptual weights of individual participants. **(C)** Comparison of behavioral confusion between tone (red) and non-tone (blue) language speakers for pitch judgments (left panel) and brightness judgments (right panel). **(D)** Relationship between years of musical training and perceptual interference of brightness on pitch judgments (green) and pitch on brightness judgments (blue). Vertical gray lines connect results obtained from the same participant. *: *p* < 0.05; **: *p* < 0.01; ***: *p* < 0.001; *n. s.*: not significant. Error bars: ±1 standard error of the mean (SEM) across participants.

Our paradigm enabled us to quantify the contribution, or perceptual weight, of the non-target dimension on the target discrimination, in terms of the slope of the linear functions shown in Fig. 1A: the steeper the slope, the more interference the non-target dimension produces on perceptual judgments (Fig. 1B; see *Data and statistical analysis* in Behavioral Experiment methods for details of the transformation). The perceptual weights assigned to the non-target dimension were significantly greater than zero for both pitch and brightness judgments (perceptual weight of brightness on pitch judgments = 0.51; two-tailed Wilcoxon signed-rank test: *N* = 20, *z* = 3.73, *p* < 0.001; perceptual weight of pitch on brightness judgments = 0.30; two-tailed Wilcoxon signed-rank test: *N* = 20, *z* = 3.10, *p* = 0.0019). The effect of irrelevant brightness changes on pitch judgments was somewhat greater than the effect of irrelevant pitch changes on brightness judgments (two-tailed Wilcoxon signed-rank test: *N* = 20, *z* = 2.02, *p* = 0.044). The positive weights indicate that participants’ discrimination was facilitated when the two features changed in the same direction and was impaired when they changed in opposite directions. The fact that the weights were significantly greater than zero confirms that participants are unable to ignore simultaneous changes in one feature (brightness or pitch) when they are judging the direction of change in the other feature (pitch or brightness), despite the feedback provided after each trial throughout the task.

Some studies suggest that sensitivity to pitch changes may differ between tone language speakers and non-tone language speakers, with one study suggesting a detrimental effect of tone-language fluency on pitch discrimination^24^ and another suggesting enhanced pitch production and discrimination in tone-language speakers^25^. We found no significant differences between tone language speakers and non-tone language speakers in terms of their DL_F0_, DL_Fc_, or the amount of perceptual interference for either pitch or brightness judgments (Mann-Whitney U-tests: *N*_tone_ = 9, *N*_nontone_ = 11; *U*_F0DL_ = 100, *p* = 0.71; *U*_FcDL_ = 90, *p* = 0.77; *U*_pitch judgment_ = 101, *p* = 0.66; *U*_brightness judgment_ = 98, *p* = 0.82; Fig. S1 and Fig. 1C). Musical training is known to facilitate pitch discrimination^26,27^, and our data are consistent with these earlier findings (Spearman’s correlation: *N* = 20; *r*_F0DL_ =-0.744, *p* < 0.001; Fig. S2). However, we found no effect of years of musical training on the perceptual interference on either pitch or brightness judgments (Spearman’s correlation: *N* = 20, *r*_pitch judgment_ = 0.184, *p* = 0.46; *r*_brightness judgment_ = 0.176, *p* = 0.33; Fig. 1D), also consistent with the findings from a previous study^11^. Because perceptual confusion does not appear to depend strongly on either musical training or linguistic background, our participants for the fMRI portion of study were selected to represent different linguistic backgrounds and a broad range of musical experience, as we would expect any neural correlates of confusion to be broadly similarly consistent across participants.

### Cortical tuning model incorporating interactions provides best description of data

We measured fMRI responses (3T, 2×2×2 mm³ resolution) from ten of the participants who had taken part in the behavioral experiment. During the scan, participants listened to sequences of short (50-ms) and long (200-ms) harmonic tones and pressed a button after each sequence to maintain alertness. The values of F0 and Fc were fixed within each sequence but varied between sequences, with each sequence sampling one out of 25 combinations of five F0 and five Fc values (see Fig. 2A and 2B and *Stimuli* in MRI Experiment methods). In an attempt to equate perceptual loudness across the stimuli, participants first completed a separate calibration task, where they adjusted the level of each of the 25 stimuli until they were perceived as equally loud (see *Procedure*). In the main experiment, these adjusted stimuli were presented 48 times each in a pseudorandom order across two experimental sessions.

**Figure 2.**
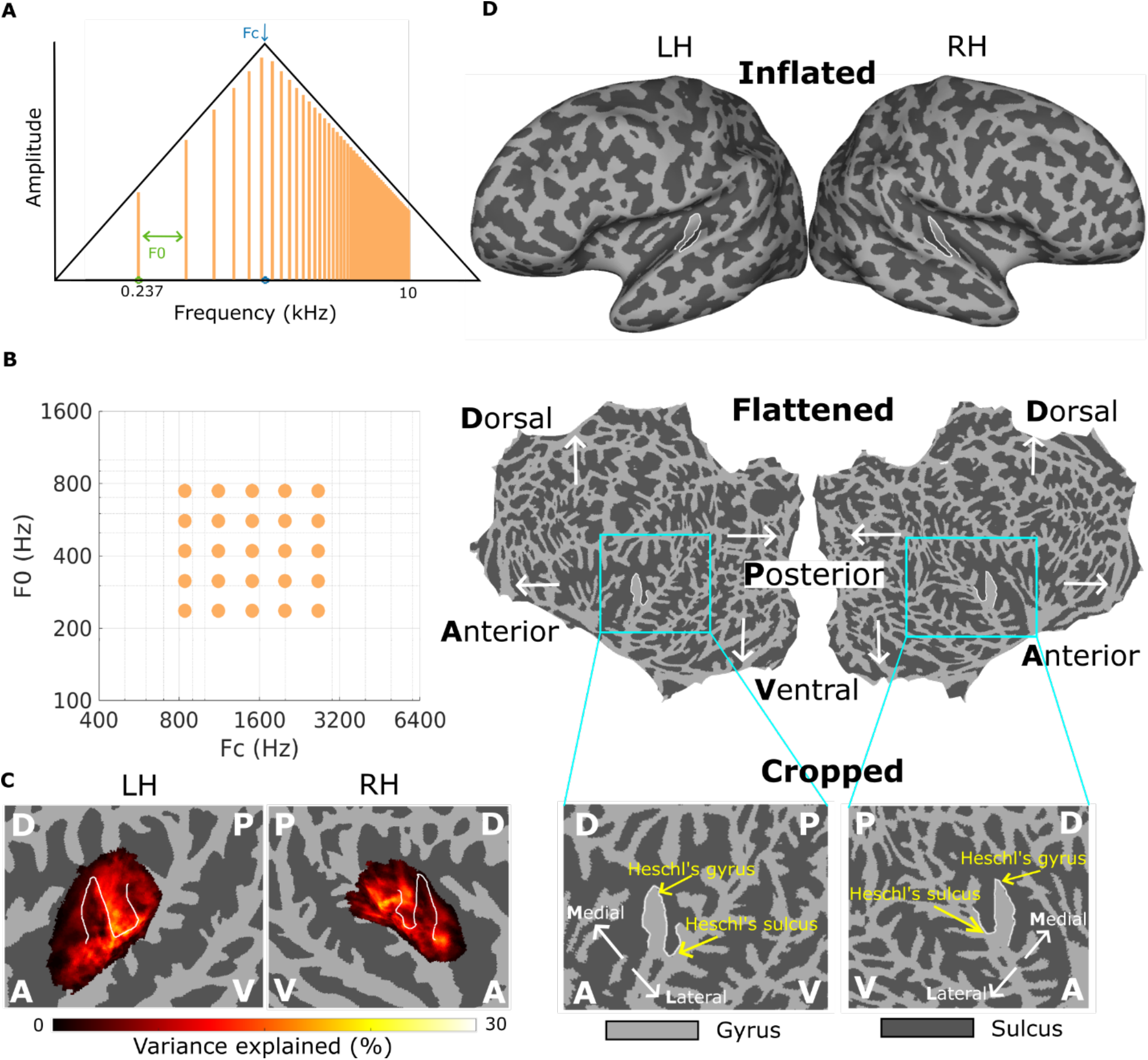
Schematic diagrams of the stimuli and visualization of the auditory cortex. **(A)** Spectrum of a harmonic complex tone used in the study plotted on log-log axes. The spacing between frequency components corresponds to the F0 (perceived as pitch). The peak of the spectrum indicates Fc (perceived as brightness). **(B)** The 25 dots represent the 25 pairs of Fc and F0 values used for the stimuli of the study. **(C)** Group-averaged variance explained by all sound stimuli against baseline silence within the group-level ROIs (regions of interest, regions that cover at least 50% of the individual ROIs, see *ROI definition*). Gyri are indicated in light grey color and sulci are indicated in dark grey color. White contours delineate the boundaries of the Heschl’s gyrus (HG) and the Heschl’s sulcus (HS). LH = left hemisphere; RH = right hemisphere; D = dorsal; V = ventral; A = anterior; P = posterior. Group average maps in the following figures all use the same plotting schemes. **(D)** Steps to prepare individual brain maps for group averaging. The inflated cortical surface (top row) is constructed from the individual participants’ anatomical T1-and T2-weighted images. The inflated brain is then flattened and mapped onto a standard fsaverage space (middle row). Two rectangular areas, one for each hemisphere (bottom row), are then specified on this standard space.

We resampled the fMRI data onto the mid-grey cortical surface via cubic interpolation^28^ (see Fig. 2D) and computed the percent BOLD change (beta value) for each trial using GLMsingle^29^. We noted that during loudness calibration, participants had assigned higher sound levels to lower-Fc tones, which may have led to over-compensation for the effects of frequency content on loudness. This created a strong correlation between BOLD responses, sound intensity, and Fc values (see correlation matrix in Fig. S3), which in turn distorted neural tuning patterns by introducing a BOLD response bias towards lower Fc values. To address this potential confound, we normalized the beta values to estimate the response to each condition at an equal sound intensity (see *Correction for the effect of sound intensity*) and used these corrected values for all subsequent analyses. We also confirmed that our key findings were replicable using the uncorrected data (Fig. S4).

To characterize the tuning properties for each vertex (the smallest unit that was imaged on the cortical surface; hereafter “voxel” for simplicity) within the regions of auditory cortex, we averaged the corrected beta values across repetitions for each of the 25 conditions. We then fitted a set of five candidate Gaussian-based encoding models to these averaged data. The models incorporated tuning to both dimensions, where the tuning in one dimension was either dependent (*interaction* model) or independent (*independent* model), incorporated tuning to just one dimension (*Fc-tuned* and *F0-tuned* models), or were responsive to sound but were not selectively tuned to either Fc or F0 (*untuned* model). These five models were hierarchically nested, with the *interaction* model mathematically subsuming all other models (see Fig. 3A and *Fitting and evaluating encoding models*).

**Figure 3.**
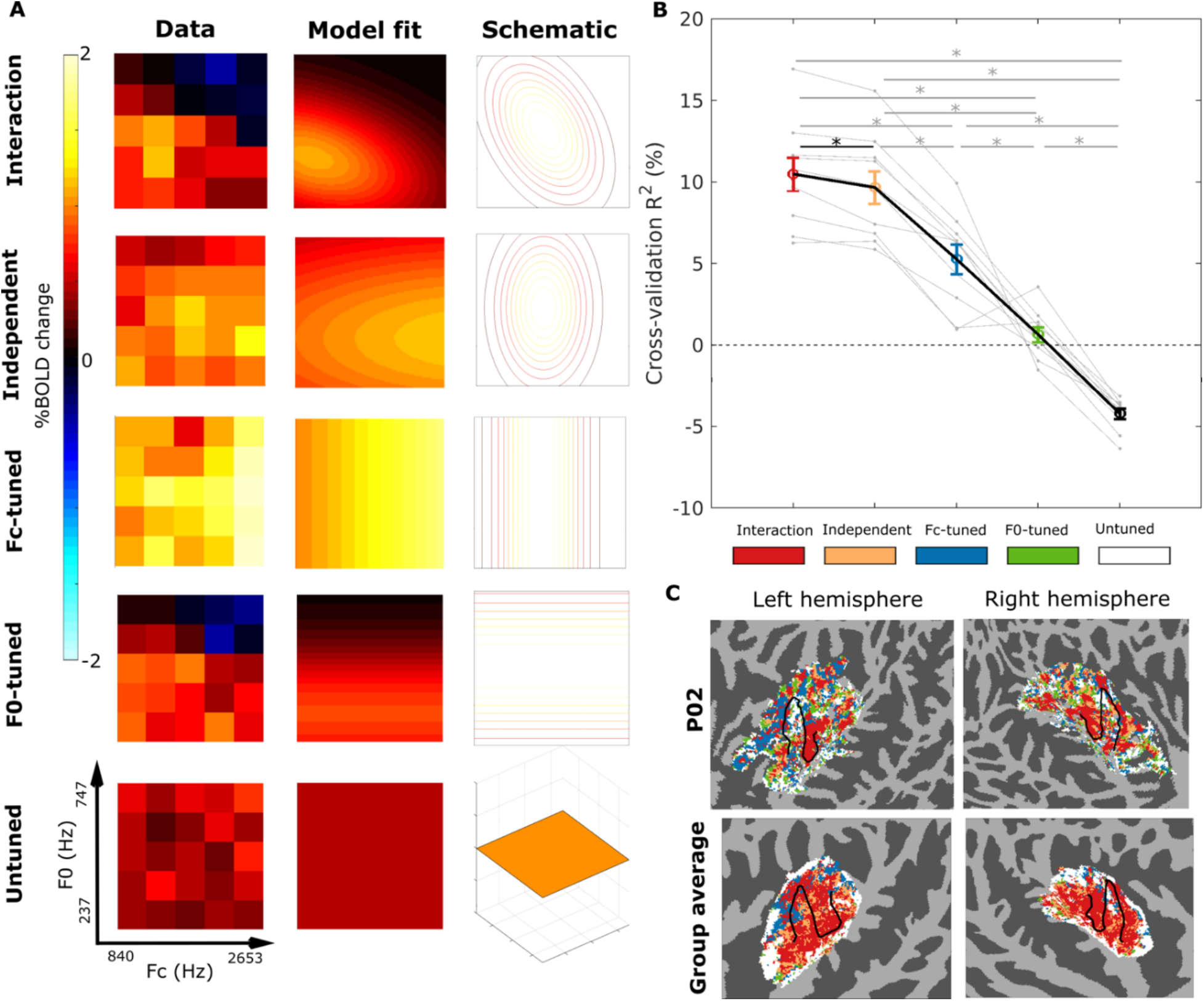
Interaction model best captures voxel-wise tuning to F0 and Fc. **(A)** BOLD responses to all 25 conditions (left column) and their model fits (middle column) of five representative voxels, one for each of the five candidate encoding models. The right column provides schematic diagrams (not reflecting real data) of the five models. The color of the matrices reflects the averaged beta value across trials. **(B)** Cross-validated R^2^ for each of the five encoding models averaged across voxels that have a positive cross-validated R^2^ for any of the encoding models except for the *untuned* model. Grey dots are individual participants’ averaged R^2^ values connected with grey lines. Circles are group means connected with black solid lines. Error bars represent ±1 SEM across participants. **(C)** Spatial distribution of voxels with different tuning properties (“best model map”). Color of the voxels within the ROI indicates the best-performing model for this voxel. Black contours delineate the boundaries of HG and HS. For the individual participant P02 (top row), only sound-responsive vertices within the ROI were shown. For the group average map (bottom row), the best model was defined as the model with the highest cross-validated R^2^, averaged across participants. *: *p* < 0.05; **: *p* < 0.01; ***: *p* < 0.001.

Figure 3A shows the cortical responses as a function of F0 and Fc from five sample voxels, each considered representative of one of the five candidate encoding models. The first and the second columns of Fig. 3A show the actual response and the predicted response by their model fit, respectively. The first voxel, well captured by the *interaction* model, shows tuning to both F0 and Fc (i.e., its responses vary to acoustic changes in both dimensions). The key property is that tuning to the two dimensions is interdependent: its preferred frequency (i.e., the peak of the fitted response) for one dimension shifts depending on the other dimension. In this case, this voxel’s preferred Fc decreases as F0 increases, resulting in larger changes in neural responses along the direction of congruent changes of Fc and F0 (from bottom left to top right) than along the direction of incongruent changes of Fc and F0 (from bottom right to top left). The direction of the interdependence in this voxel aligns with the perceptual results and with predictions of efficient neural coding, which posit that sensory systems maximize discriminability for statistically common stimulus variations^16,17,30^ — in this case, the positive correlation between Fc and F0^14^.

The second voxel, captured by the *independent* model, is also tuned to both dimensions, but its preferred frequency for each is fixed and does not depend on the other. The third and fourth voxels exemplify tuning to a single dimension – either Fc (*Fc-tuned*) or F0 (*F0-tuned)*, respectively – but invariance to the other dimension. The fifth voxel responds to sound but shows no clear tuning preference for either dimension (*untuned*).

Across the entire auditory cortex, models featuring tuning to both dimensions significantly outperformed those tuned to one or none (Fig. 3B). The *interaction* model held a small but significant advantage over the *independent* model (Friedman test across five models: c^2^(4) = 38, *p* <.001; post-hoc pair-wise two-tailed Wilcoxon sign-rank test between the *interaction* model and the *independent* model: *N* = 10, *W* = 54, Bonferroni corrected *p* =.04). Importantly, this superior performance was not an artifact of greater model complexity, as we used a leave-one-run-out cross-validation procedure that controls for overfitting (see *Fitting and evaluating encoding models*). There were also significant differences in performance between all other pairs of conditions (Bonferroni corrected *p* < 0.05 in all cases).

To visualize the spatial distribution of voxels with different tuning properties, we plotted the best model (the one that produced the best cross-validated prediction among the five models) of each voxel within the region of interest (ROI, see *ROI definition*) on the cortical surface (Fig. 3C). Voxels best described by the *interaction* and the *independent* models spanned the Heschl’s gyrus (HG) and the Heschl’s sulcus (HS), whereas voxels best fit by single-dimension models (*Fc-tuned* or *F0-tuned)* were sparser and tended to flank these regions. Comparing the two single-dimension models, the *Fc-tuned* model performed better (Fig. 3B) and provided the best fit to more voxels than the *F0-tuned* model (Fig. 3C; see Fig. S5 for individual maps). This suggests that tuning to Fc is more robust and more widespread than tuning to F0 within auditory cortex, consistent with earlier findings of robust tonotopy across auditory cortex^31^.

Taken together, these results demonstrate that the model incorporating the interaction between tuning to pitch and brightness best captures the tuning patterns of the auditory cortex, a property that can support efficient coding of the natural covariation of the two features. Given its superior model performance and its flexibility (as it mathematically subsumes all other candidates, see *Fitting and evaluating encoding models*), we used the predicted beta weights from the *interaction* model for all subsequent analyses of local and global tuning patterns.

### Within-voxel substrates of perceptual confusion

Having established the existence of “interaction voxels” with tuning dependence of one dimension on the other, we next investigated whether the direction of the tuning dependence is consistent with the perceptual confusion and the predictions of efficient coding. To answer this question, we looked at the shifts in preferred F0 across different values of stimulus Fc and vice versa. When comparing the preferred Fc across low and high stimulus F0s, we observed a “warmer” map of preferred Fc values as the stimulus F0 increased from low to medium to high (**Error! Reference source not found.**A), indicating that voxels’ preferred Fc tend to decrease with increasing stimulus F0. This trend is illustrated in the *difference* map of preferred Fc between the highest and lowest F0 (High–Low; bottom row in **Error! Reference source not found.**A), which exhibits cool colors overall, suggesting a downward shift in the preferred Fc as the F0 increases. The F0 preference map showed a similar pattern (**Error! Reference source not found.**B), with preferred F0 decreasing with increasing stimulus Fc. Quantifying this effect, we found that as the stimulus F0 increased from lowest to highest, significantly more voxels decreased their preferred Fc than increased it (two-tailed Wilcoxon signed-rank test: *N* = 10, *W* = 49, *p* = 0.027); similarly, as the stimulus Fc increased from lowest to highest, significantly more voxels decreased than increased their preferred F0 (two-tailed Wilcoxon signed-rank test: *N* = 10, *W* = 54, *p* = 0.004). This effect was highly robust and can be seen in individual participants (Fig. 4D and Table S1). Additionally, the difference between the number of voxels shifting downwards and upwards in preferred F0 was significantly greater than that of preferred Fc (two-tailed Wilcoxon signed-rank test: *N* = 10, *W* = 55, *p* = 0.002). In other words, brightness changes had a greater effect on the preferred pitch than pitch changes had on the preferred brightness, consistent with our perceptual findings showing a stronger non-target weighting of brightness on pitch judgments than the reverse (Fig. 1B).

**Figure 4.**
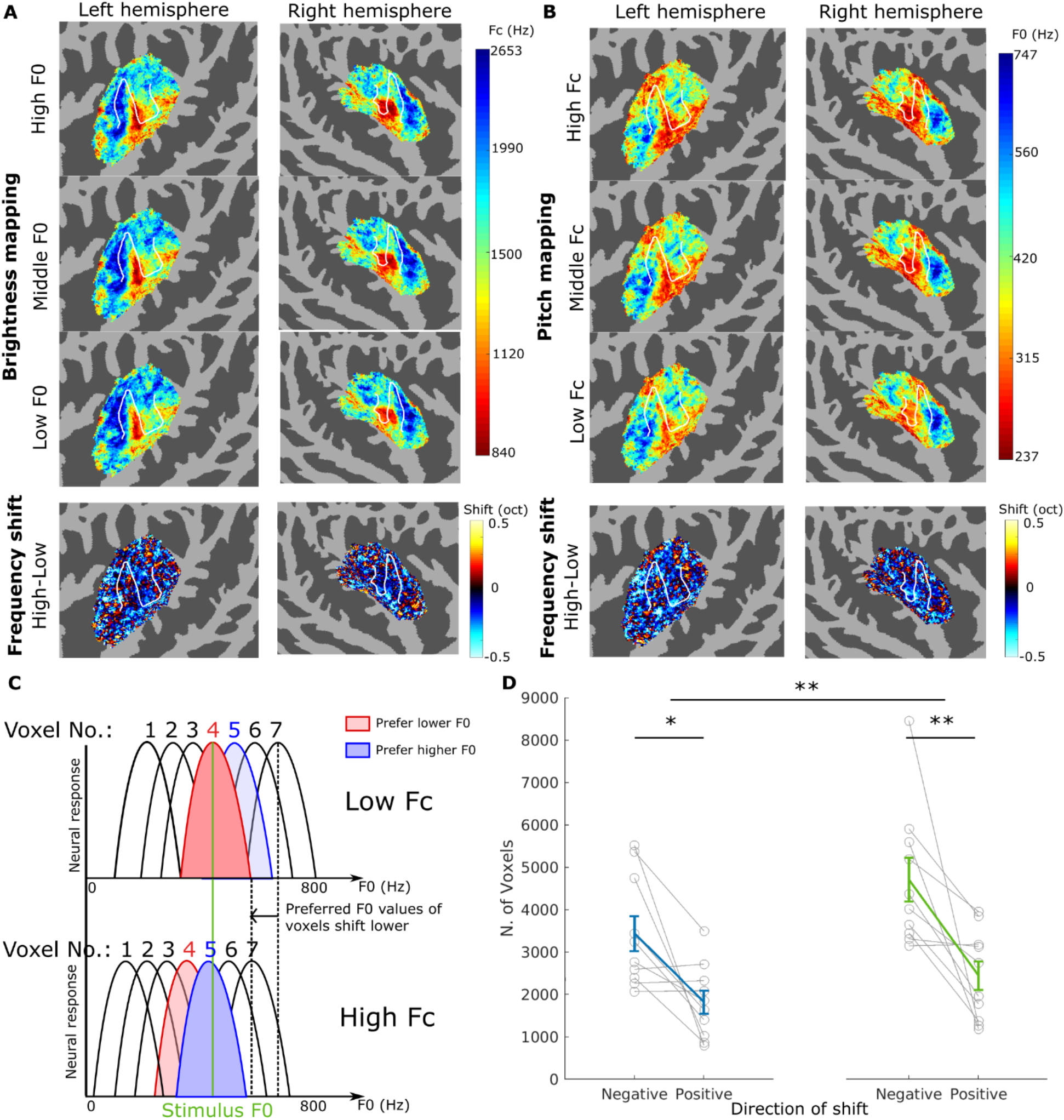
Within-voxel interaction between F0 and Fc as a potential substrate of perceptual confusion. **(A)** Top three rows: group average maps of preferred Fc values at the lowest (237 Hz), middle (420 Hz), and the highest (747 Hz) F0s. Bottom row: difference in preferred Fc values between the highest and the lowest F0s. **(B)** Top three rows: group average maps of preferred F0s at the lowest (840 Hz), middle (1500 Hz), and the highest (2653 Hz) Fc values. Bottom row: difference in preferred F0s between the highest and the lowest Fc values. Group averaging and subtraction of preferred frequencies were computed on a log scale. **(C)** Schematic diagram explaining how within-voxel interaction between F0 and Fc may underlie perceptual confusion. Top row: neural response of voxels as a function of F0 at a low Fc. Bottom row neural response of the same group of voxels as a function of F0 at a high Fc. The example stimulus indicated by the green line excites Voxel 4 (shaded in red to indicate a relatively lower preferred F0) the most at low Fc but excites Voxel 5 (shaded in blue to indicate a relatively higher preferred F0) the most at high Fc, implying a higher perceived pitch. (D) Left: number of voxels with a positive shift or a negative shift in preferred Fc from a low stimulus F0 to a high stimulus F0. Right: number of voxels with a positive shift or a negative shift in preferred F0 from a low stimulus Fc to a high stimulus Fc.

To illustrate how the direction of this downward preferred frequency shift is consistent with perceptual confusion, the schematic in Fig. 4C depicts the neural response level as a function of F0 from an array of voxels at low versus high Fc values. Assuming that the perceived pitch and brightness of a stimulus are indicated by the voxels with the strongest response to the stimulus, if a given F0 of the stimulus (green vertical line) leads to maximal stimulation of one voxel (Voxel 4 in Fig. 4C) at a low Fc, then at a higher Fc, when all the voxels’ preferred F0s shift downward, the same stimulus F0 will lead to maximal stimulation of a voxel that is tuned to a higher F0 than Voxel 4 (Voxel 5 in Fig. 4C), resulting in the perception of a higher pitch. Similarly, if a given stimulus Fc leads to maximal stimulation of one voxel at a low F0, the same Fc will lead to maximal stimulation of a voxel that is tuned to a higher Fc, resulting in the perception of a brighter timbre.

In summary, we observed the neural tuning dependency, or representational interaction between pitch and brightness within individual voxels in the same direction as predicted by the well-known perceptual confusion effects between pitch and brightness. We also found stronger voxel tuning dependence of preferred pitch on brightness than the dependence of preferred brightness on pitch, also consistent with our perceptual findings. Both the voxel tuning dependency and the asymmetry between the pitch dependence on brightness and the brightness dependence on pitch were robustly observed at the individual and group levels, in line with the perceptual effects.

### Global cortical substrates of perceptual confusion

One of the most established tuning properties throughout the auditory pathways, from the cochlear to the auditory cortex, is the orderly frequency-to-place mapping, or tonotopy^32,33^. Such orderly mapping has also been found for Fc and, to some extent, F0 in the auditory cortex^31^. To examine if such orderly mapping of Fc and F0 exists in our data, we show the preferred Fc value at the middle F0 (420 Hz) for each voxel (top row, Fig. 5) and the preferred F0 value at the middle Fc (1500 Hz) for each voxel (bottom row, Fig. 5), both based on the fitted responses from the *interaction* model. Interestingly, the two maps show many similarities, with both exhibiting high-low-high gradients that are spatially overlapped. Specifically, low-frequency-preferred regions for both mappings are in the anterolateral HG, with both also showing two low-high gradients stemming from the same low-frequency-preferred region and extending out in the anteromedial and posterior directions. Such a pattern is evident in individual participants (Fig. S6) and is also reflected in the significant positive correlation between the preferred Fc and preferred F0 across the auditory cortex (Spearman’s correlation, ρ > 0.22, *p* < 0.001, for each of the 10 participants). The overlapping gradients imply that increases (or decreases) in Fc lead to changes in activation patterns that are similar to those resulting from increases (or decreases) in F0. For example, increases in Fc alone and in F0 alone would both lead to a decrease of response in the anterolateral HG (low-frequency-preferred region) and an increase of response outwards (high-frequency-preferred regions) (magenta arrows in Fig. 5). Such similarity provides a global cortical substrate of perceptual confusion between pitch and brightness. Note that this *global* substrate is distinct from the voxel-level findings in the previous section. The previous substrate is derived from the tuning dependency or interaction between the two features *within* individual voxels, a property that supports the efficient joint encoding of pitch and brightness at the local population level, whereas the current substrate is derived from the spatial relationship *between* voxels with different preferred F0 and Fc values, a correlate of the pitch-brightness confusion at the global scale across the auditory cortex. Nevertheless, they are both consistent with the behavioral evidence, showing that a change in one feature (pitch or brightness) can be confused for a change in the same direction of the other.

**Figure 5.**
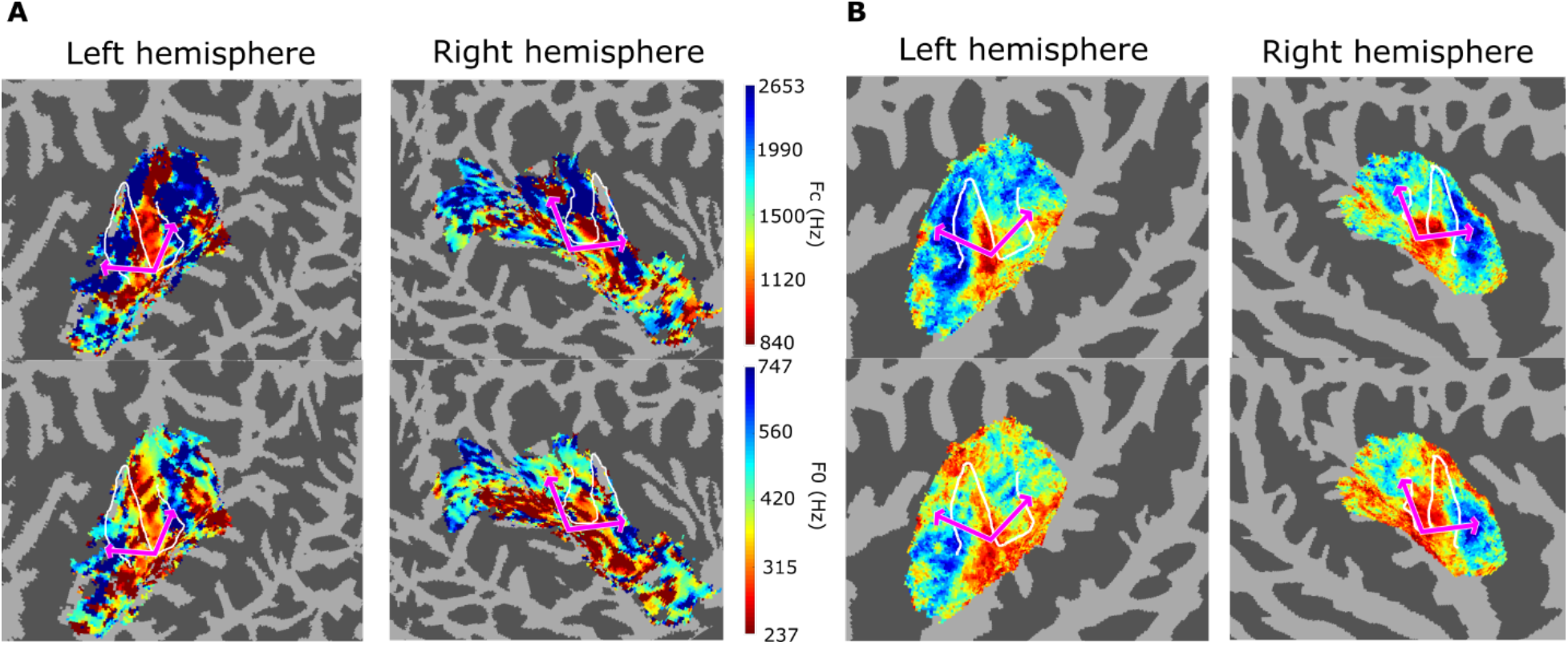
Pitch and brightness mappings have overlapping gradients. **(A)** Pitch and brightness mappings for an individual participant (P07). **(B)** Pitch and brightness mappings averaged across 10 participants. Top row: preferred Fc at the middle F0 (420 Hz). Bottom row: preferred F0 at the middle Fc (1500 Hz). Magenta arrows indicate the trajectories going from a low-frequency-preferred region to a high-frequency-preferred region for both F0 and Fc at the same locations on the cortex.

### Combining within-and across-voxel patterns to predict perceptual confusion

We next sought to determine if the perceptual confusion between pitch and brightness is reflected in the neural activation pattern across the auditory cortex. Our behavioral results showed that stimulus pairs with congruent changes in pitch and brightness were easier to discriminate than pairs with incongruent changes. The efficient coding principle also predicts that neural populations increase discriminability along the expected, or frequently present, covariations^16,17,30^, producing more distinct neural responses when pitch and brightness change in the same direction than in opposite directions. Based on this principle, we hypothesized that pairs of conditions with congruent changes in F0 and Fc would elicit more distinct (i.e., less similar) neural activation patterns than incongruent pairs.

To test this hypothesis, we computed the cosine similarity between activation patterns across the auditory cortex of every pair of stimulus conditions. We then organized these values into a representational similarity matrix, structured by the difference in Fc (DFc) and the difference in F0 (DF0) between pairs of stimuli, with positive differences indicating congruent changes (e.g., an increase in both F0 and Fc) and negative differences indicating incongruent changes (e.g., an increase in F0 and a decrease in Fc) within each pair. In the similarity matrix (Fig. 6), we observed lower similarity values in the upper left and lower right of the matrix compared to the lower left and upper right of the matrix, which indicated that pairs of conditions with congruent changes have lower neural similarity than pairs with incongruent changes. This pattern, which is evident in both the group-average matrix (Fig. 6) and in the data from all ten individual participants (Fig. S7), shows that the fMRI representations of congruent changes in pitch and brightness are more discriminable than incongruent changes, in line with our perceptual results, and consistent with expectations based on efficient coding principles. It is important to note that both the voxel-based tuning dependencies and the global topography of preferred F0 and Fc contribute to similarity matrices (see simulation results in Fig. S8), and thus serve as separate potential substrates for perceptual confusion.

**Figure 6.**
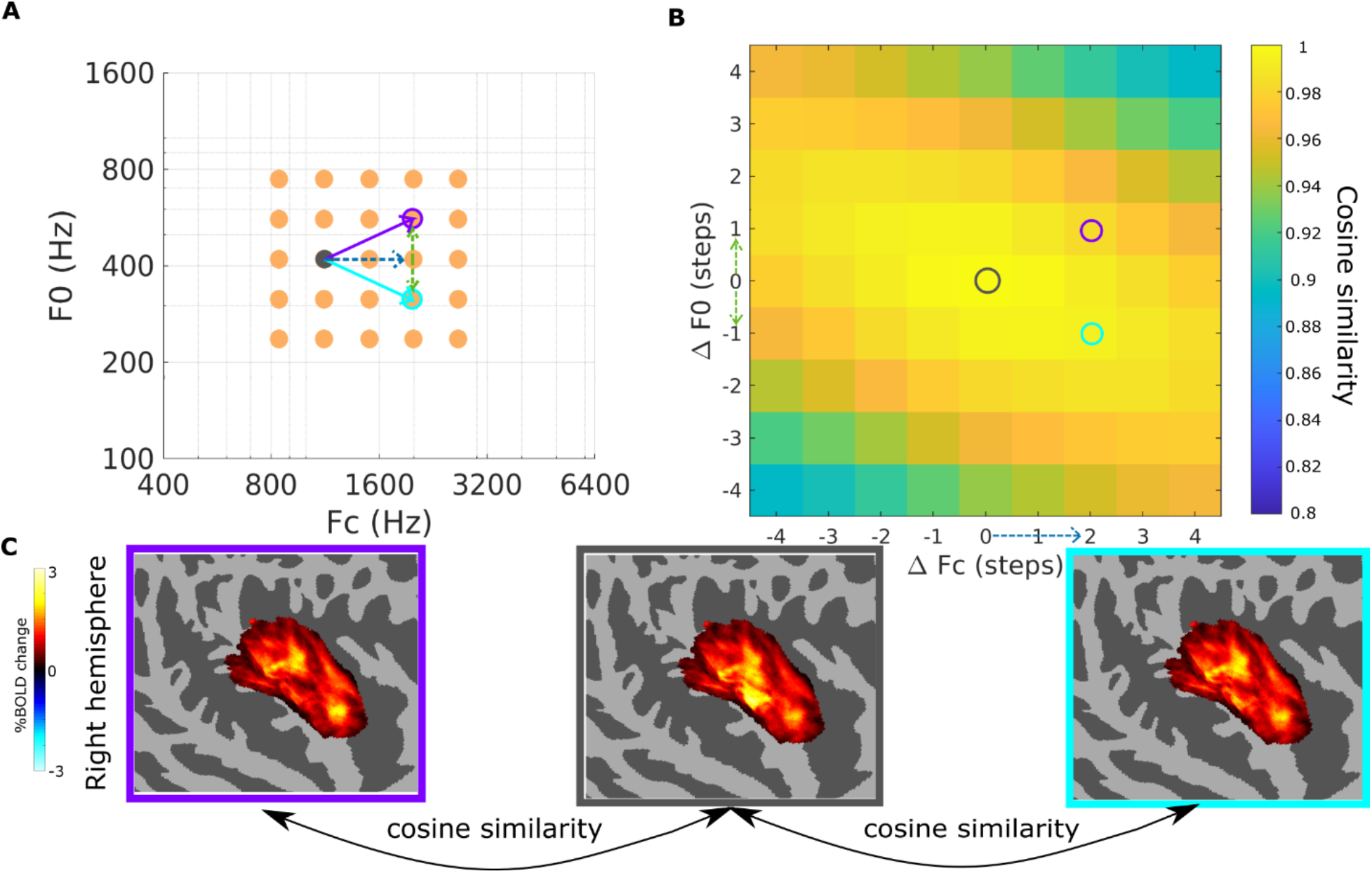
Cortical activation patterns are less similar for congruent changes than for incongruent changes between pitch and timbral brightness. **(A)** Stimulus grid showing two example pairs of conditions. The congruent pair of conditions (grey and purple) are two steps away in Fc (blue dashed arrow) and one step away in F0 (green dashed arrow) with Fc and F0 both increasing / decreasing. The incongruent pair of conditions (grey and cyan) are two steps away in Fc (blue dashed arrow) and one step away in F0 (green dashed arrow) with Fc and F0 changing in opposite directions. **(B)** Similarity matrix showing cosine similarity between pairs of conditions. Similarity of the congruent pair is placed at position [2, 1] (purple circle) and similarity of the incongruent pair is placed at position [2,-1] (cyan circle) along with all other pairs with the same set of differences in F0 and Fc. Cosine similarity values were averaged across all pairs of conditions populating the same location of the matrix. **(C)** Cortical activation patterns of the three conditions. Purple = condition with congruent change from reference; grey: reference; cyan: condition with incongruent change from reference.

### Comparing hemispheric tuning to pitch and brightness

When we compared the overall tuning to *both* F0 and Fc, reflected in the performance of the *interaction* model, the number of voxels favoring the *interaction* model did not differ significantly between hemispheres (two-tailed Wilcoxon signed-rank test: *N* = 10, *W* = 19, *p =* 0.432), even though overall performance of the *interaction* model was marginally better for the right hemisphere than for the left (two-tailed Wilcoxon signed-rank test: *N* = 10, *W* = 5, *p =* 0.020).

We also compared hemispheres in their *exclusive* tuning to F0 or Fc separately by examining the performance of the *F0-tuned* and the *Fc-tuned* models. The numbers of voxels favoring the *F0-tuned* model did not differ significantly between hemispheres (two-tailed Wilcoxon signed-rank test: *N* = 10, *W* = 18.5, *p =* 0.375); the same was true for the *Fc-tuned* model (two-tailed Wilcoxon signed-rank test: *N* = 10, *W* = 28, *p =* 1). Again, however, in terms of model performance, the right hemisphere showed marginally better performance than the left for the *Fc-tuned* model (two-tailed Wilcoxon signed-rank test: *N* = 10, *W* = 5, *p =* 0.020), but not for the *F0-tuned* model (two-tailed Wilcoxon signed-rank test: *N* = 10, *W* = 18, *p =* 0.375). In sum, we did not observe a hemispheric difference in terms of the *size* of the cortical regions tuned to these two features, and any small differences in coding accuracy were only marginally significant.

## DISCUSSION

In this study we investigated the neural basis for the long-recognized perceptual phenomenon of pitch– brightness confusion, where changes in one dimension are often mistaken for changes in the other. Some studies have hypothesized that this confusion occurs at the encoding level, reflecting an efficient coding strategy that adapts to correlated pitch and brightness in natural sounds^17,18,30^. However, others have instead suggested that confusion arises at a later decision stage^34^, comparable to the well-known Stroop interference effect^35^. Behavioral measures alone have struggled to distinguish between these two hypotheses. We used fMRI-based measures to test for evidence of encoding-level interactions between pitch and brightness in a paradigm that required no decisions or judgments of pitch or brightness. Our findings provide two distinct lines of evidence from human auditory cortex—at both local and global scales— that suggest interactions between pitch and brightness at the encoding level.

### Local interactions between pitch and brightness reflect efficient coding

Our first, and most critical, finding is the interaction, or tuning dependency between pitch and brightness tuning at the local, voxel-wise level. Leveraging the fully crossed stimulus design, we demonstrate that the cortical representation of pitch or brightness can be characterized as a tilted Gaussian function of stimulus F0 and Fc at the encoding level, mirroring our behavioral results showing performance for directional judgment of one dimension as the weighted sum of variations in both dimensions (Fig. 1A-B). Crucially, we observed that the preferred pitch of individual voxels systematically decreased as brightness increased (and vice versa). This tuning pattern (see a representative depiction in the first row of Fig. 3A) exhibits steeper changes—and thus better discriminability—along the direction of congruent changes between pitch and brightness compared to incongruent changes and matches the direction of perceptual confusion^8–11^.

Interestingly, the observed pattern of the representational interaction aligns with the predictions of efficient coding. This theory posits that sensory processing is designed to reduce information redundancy^15^ and maximize the efficiency of stimulus representation by allocating neural resources based on statistical regularities in the environment^16^. In natural sounds, such as those produced by the human voice and by musical instruments, F0 and Fc are often positively correlated^13,14^. An efficient system should therefore increase discriminability along this “expected” direction of co-variation through exposure to these regularities^15–17^, leading to better discrimination with the congruent changes in pitch and brightness than incongruent changes. This interpretation is further supported by a developmental study showing that pitch discrimination and brightness discrimination of infants—who have had relatively less exposure to these natural statistics—were less susceptible to changes in the other dimension (i.e., less confusion between pitch and brightness) than adult non-musicians^18^. Thus, our finding of steeper neural response changes along congruent variations of pitch and brightness than incongruent ones demonstrates a neural implementation of efficient coding.

It is important to note, however, that our data also revealed voxels that were tuned to both pitch and brightness, but where the tuning of one feature was unaffected by the level of the other (*independent* model, see Fig. 3). The existence of these voxels is consistent with our perceptual experience that pitch and brightness are not always collapsed onto a unitary dimension. For instance, pitch-matching experiments have shown that listeners can accurately match the frequency of a pure tone to the F0 of a complex tone across different levels of brightness or spectral content^36^. The coexistence of both interaction and independent coding may reflect a flexible, task-dependent encoding scheme: it preserves separable information of pitch and brightness for feature fidelity, while also capturing the statistical regularities of the natural world when detecting and following variations in both features over time.

### Global gradients and population-level discriminability of pitch and brightness changes

Complementing the local, voxel-wise interaction, our second major finding is that pitch and brightness share a common global tuning gradient across the auditory cortex. This implies that an increase in pitch and an increase in brightness elicit cortical activation that follow a shared spatial trajectory. This finding extends the previous findings of co-localized pitch-and brightness-sensitive neural populations^20–22,37^ from a local perspective to a global property of the auditory cortex, demonstrating that these neural populations are not only tuned to both features, but also topographically organized along a shared axis across the auditory cortex. This shared global map provides a macro-scale cortical substrate for the confusion, organizing the co-varying features along a single, efficient gradient of cortical activation.

Finally, we sought to determine how the local interaction and the global gradients combine to shape the overall representation property of the auditory cortex. By analyzing the activation patterns of the entire auditory cortex, incorporating both local tuning shifts and global distributions, we observed lower similarity (and thus better discriminability) for congruent changes in pitch and brightness compared to incongruent changes. This observation confirms that the principles of efficient coding are not restricted to isolated voxels but are a pervasive feature across the auditory cortex. Together, these local and global representation patterns provide a comprehensive neural explanation for why pitch and brightness are perceptually confused, grounding this well-known phenomenon in the fundamental principles of efficient sensory encoding.

### Anatomical consistency with prior work

The spatial distributions of pitch-tuned and brightness-tuned voxels in our results are in good agreement with previous findings that investigated each dimension separately^31,33,38^. In Heschl’s gyrus (HG) and Heschl’s sulcus (HS), where joint tuning to pitch and brightness was mostly likely to be observed (Fig. 3C, red and orange voxels), the V-shaped high-low-high gradient matched the location and spatial layout of pure-tone tonotopy^33,39^ and tuning to spectral peaks of complex tones^31^. In the surrounding regions, where exclusive tuning to pitch or brightness was more likely to be observed (Fig. 3C, green and blue voxels), brightness mapping (top row of Fig. S9A) still resembled tonotopy, showing a high-low-high gradient spanning HG, as expected based on previous results^31^. Pitch mapping, on the other hand, formed a high-low-high gradient *surrounding* HG (middle row of Fig. S9A). Such a pattern aligns well with findings from an earlier study of just Fc or F0 tuning^31^ (bottom row of Fig. S9A). In addition, pitch mapping can be seen as having a low-high-low gradient along the long axis of HG (Fig. S9B). In terms of relative strengths of tuning, both our results (Fig. S10) and a previous study^31^ showed stronger brightness tuning than pitch tuning, especially in the core auditory regions, but somewhat stronger pitch tuning than brightness tuning in the surrounding regions (see group average map in Fig. S10 and individual maps in Fig. S11). These results not only replicate the previous findings but also delineate a clearer picture of pitch and brightness mapping in the human auditory cortex, with joint tuning in HG and HS and distinct gradients in the surrounding regions.

### Sharper transitions of brightness mapping compared with pitch mapping

The brightness mappings had sharper transitions between low-and high-frequency-preferred regions than the pitch mappings (Fig. 5) on the anterior border of HG. A potential explanation comes from the relationship between the stimulus frequency range and the effective frequency ranges for pitch and brightness perception in human listeners. The range of F0s for which listeners can hear a clear melodic pitch extends from roughly 30 to 4,000 Hz, corresponding to about 7 octaves^42–44^, whereas the range of audible frequencies, and hence the range of potential Fc values, for young listeners with normal hearing has a much wider range of 20 to 20,000 Hz, or about 10 octaves. Yet, the range of the stimuli for Fc and F0 in this study was the same, meaning that the proportion of the effective frequency range that our stimuli covered was smaller for Fc than for F0. Assuming a uniform representation along the frequency axis, we would expect the frequency range of our stimuli to encompass a smaller proportion of the range of voxels sensitive to Fc than to F0, meaning that Fc tuning curves are more likely to appear as a slope (with the preferred Fc at the low or high edge of the sampled frequency range) than a peak (with the preferred Fc within the sampled frequency range). This difference in range may explain why the brightness map in this study appears to be more binarized than the pitch map.

### Hemispheric differences

There are debates on whether hemispheric differences exist in the cortical representation of pitch, timbre, and other salient spectral and temporal dimensions^45,46^. The current study found evidence for a small right hemisphere advantage in the cortical *tuning* to pitch and brightness but no evidence for asymmetry in the *size* of cortical regions exhibiting tuning to these two features. For representations of pitch, while bilateral activation is observed in almost all studies with comparable sizes of activated regions between hemispheres^31,47^, the right hemisphere has been found to have a small advantage in the magnitude of activation or selectivity, usually in the cases of pitch changes or melodies^45,48–50^. On the other hand, cortical representations of pitch salience, which is usually obtained with a fixed pitch, have generally demonstrated hemispheric symmetry^47,51,52^. For representations of brightness, variations in spectral envelope also elicit bilateral activation^21,37^, although a non-significant trend of right lateralization in the sensitivity to spectral variation has been reported^37^. Thus, the findings of the current study, which little or no hemispheric differences in encoding of pitch or timbral brightness, are in general agreement with most but not all previous findings. Differences may be due to the nature of the stimuli, in terms of whether longer melodies or simple repeated tones were used.

### Consistency of effects across individuals

Our experiments focus on within-subject effects between conditions, embracing an approach of ‘deeper’ studies (i.e., more repetitions) on fewer participants, which has proved successful in other neuroimaging studies^53^. Our approach receives support from the observation that both our perceptual and neural findings were robust across participants. In the perceptual results with 20 participants, all but three demonstrated confusion in both pitch and brightness judgments in the expected direction (positive perceptual weights of the irrelevant feature; see Fig. 1B). In the neural results, although ten participants are too few to explore brain-behavior correlations, all ten participants showed within-voxel tuning dependence between pitch and brightness in the expected direction (more negatively shifting voxels than positively shifting voxels; see Fig. 4D), and showed substantial within-participant correlations between pitch mapping and brightness mapping (see Fig. S6). The results at the single-voxel and population levels are further illustrated in the consistency found in the tilting direction of the similarity matrices across all ten participants (see Fig. S7), consistent with the direction of the perceptual interference. Our study was not designed to test for effects of factors such as musicianship and tonal language background. Nevertheless, in our limited sample, interactions between pitch and timbral brightness appear to be robust across these factors, both at perceptual and neural levels.

### Limitations of the study

One caveat pertaining to all fMRI studies is that voxel responses reflect locally pooled neural activity within the volume of a voxel. In our case, joint encoding of pitch and brightness at the voxel level could in principle reflect different neuronal populations within the voxel, each uniquely encoding one of those dimensions. Along similar lines, voxels with unique selectivity to pitch or timbre may reflect a greater concentration of neurons selective to one or other dimension, rather than unique selectivity. Evidence from single-unit recordings in ferret suggest that it is in fact common for individual neurons to be selective for both pitch and timbre, as well as spatial location^20^. Although that study was not designed to explore interactions between dimensions, it provides evidence that individual neurons can indeed be selective for multiple dimensions.

## RESOURCE AVAILABILITY

### Lead contact

- Requests for further information and resources should be directed to and will be fulfilled by the lead contact, Andrew J. Oxenham.

### Materials availability

- This study did not generate new unique reagents.

### Data and code availability

- Deidentified data reported in this paper will be shared by the lead contact upon reasonable request.
- The original code for the behavioral and fMRI data analysis and experimental control will be made available by the lead contact upon reasonable request.
- Any additional information required to reanalyze the data reported in this paper is available from the lead contact upon request.

## ACKNOWLEDGMENTS

This work was supported by NIH grant R01 DC005216 (awarded to AJO). We thank Jaeeun Lee for assistance in data collection and Juraj Mesik for helpful advice.

## AUTHOR CONTRIBUTIONS

Conceptualization, A. J. O.; methodology, E. J. A., K. N. K., Y. O., A. J. O.; investigation, E. J. A., Y. O.; writing – original draft, Y. O., A. J. O.; writing – review & editing, E. J. A., K. N. K., Y. O., A. J. O.; funding acquisition, A. J. O.; resources, K. N. K., A. J. O.; supervision, K. N. K., A. J. O.

## DECLARATION OF GENERATIVE AI AND AI-ASSISTED TECHNOLOGIES IN THE WRITING PROCESS

During the preparation of this work, no generative AI or AI-assisted technologies were employed, beyond the automatic spelling and grammar checking incorporated within Microsoft Word. The authors take full responsibility for the content of the publication.

## SUPPLEMENTAL INFORMATION

**Document S1.** Figures S1–S14, Tables S1-S3.

---

**Table 1.**
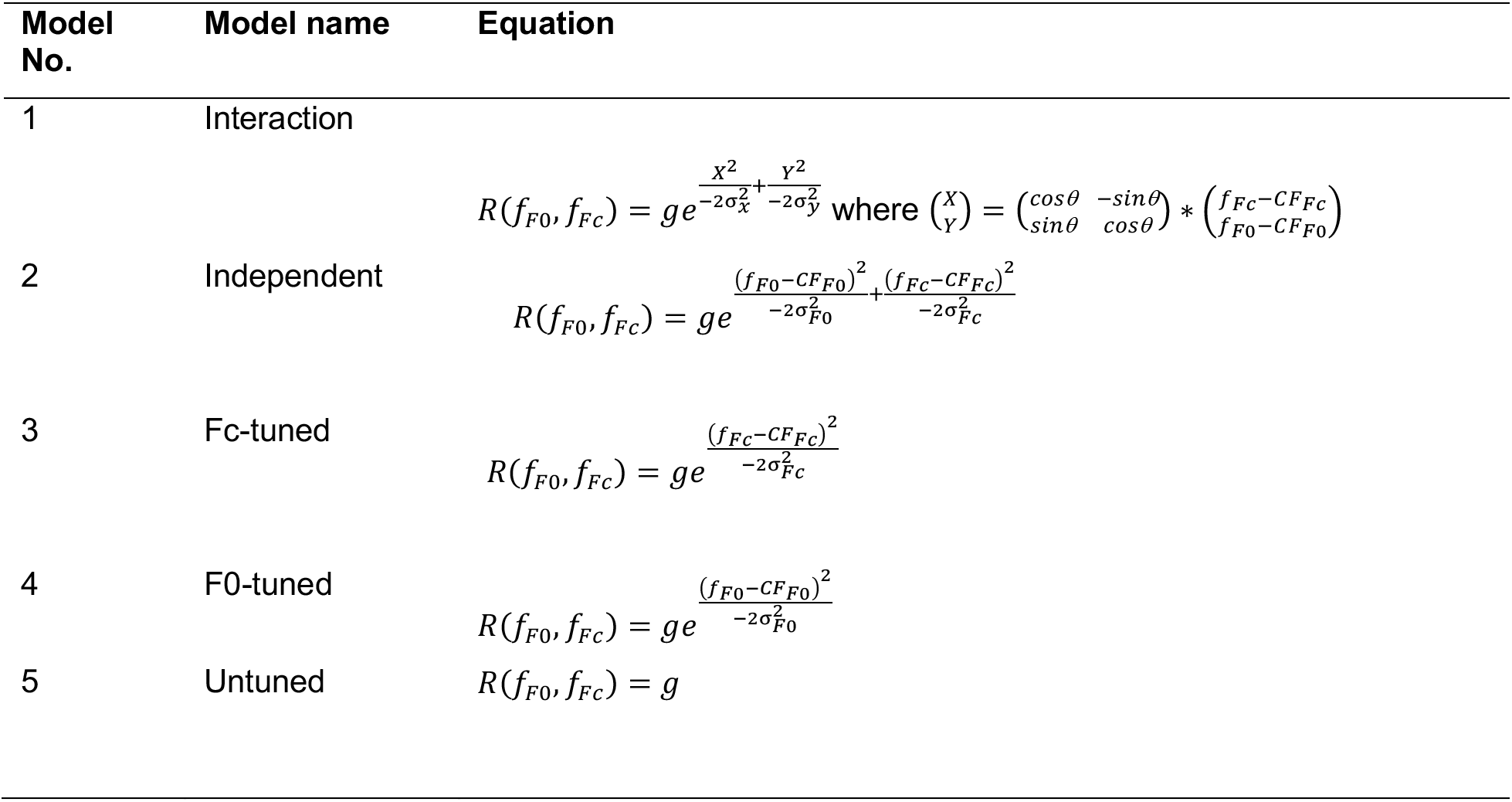
Equations of the five encoding models.

## STAR★METHODS

### METHOD DETAILS

#### MRI Experiment

##### Participants

Ten participants [five females, four males, one non-binary, mean (SD) age of 30.9 (9.3) years, ranging from 23 to 53 years, all right-handed] were recruited from the University of Minnesota community. Three of the participants were authors (Y. O., E. J. A., and A. J. O.). All participants had normal hearing (pure-tone thresholds no higher than 20 dB hearing level (HL), at octave frequencies between 250 Hz and 8 kHz) except for one who had a slightly elevated pure-tone threshold of 25 dB HL at 4 kHz in their right ear. One participant reported having intermittent tinnitus. Five of the ten participants were identified as non-musicians, having two or fewer years of formal musical training. The other five participants had an average of 12.4 years of musical training on non-percussion instruments. Three of the ten participants were tone-language speakers (Mandarin Chinese or Cantonese). Years of musical training and language background of individual participants are reported in Table S2. The experimental protocol was approved by the University of Minnesota Institutional Review Board (Approval No.: 1207M16625). All participants provided written informed consent before the experiment.

##### Stimuli

Stimuli were generated digitally in MATLAB (The MathWorks) and presented with its Psychophysics Toolbox extensions (*53*). The stimulus set consisted of 25 harmonic complex tones, each of which was bandpass filtered with 12 dB/oct slopes, then filtered with a 16^th^-order lowpass filter and a cutoff frequency of 10 kHz (see Fig. 1A). Tones varied both in F0 (between 237 and 747 Hz), affecting pitch, and in the bandpass filter’s center frequency (Fc; between 840 and 2653 Hz), affecting brightness, in five equal steps (on a logarithmic scale) for both dimensions (i.e., F0 = 237, 315, 420, 560, or 747 Hz; Fc = 840, 1120, 1500, 1990, or 2653 Hz). Each F0 was paired with each Fc in a 5×5 design, for a total of 25 unique stimuli (see Fig. 1B).

The ranges of F0 and Fc were selected to ensure that the F0 was always below the Fc (allowing the Fc to be defined by the amplitudes of the harmonic components) and, critically, that all stimuli contained a sufficient number of spectrally resolved harmonics to elicit a salient pitch. Harmonics are typically considered resolved up to the 10th^54^. Our stimuli always had between six and ten of the first ten harmonics audible (within 30 dB of peak amplitude), and 16 of the 25 conditions had all ten audible. Pitch salience was also confirmed by measuring listeners’ sensitivity to F0 and Fc changes at the four corners of the stimulus matrix, as well as at the center; see Fig. S12) to rule out potential confounds from harmonic resolvability.

The number of steps in F0 and Fc was based on data quality considerations (each condition presented 24 times per session) and the total duration of the experiment (two hours for each of the two sessions). Given these constraints, we determined that five levels per dimension, totaling 25 conditions, provided a good balance between the sampling density of the feature space and the total duration of the experiment.

##### Procedure

Stimuli were presented through MR-compatible Sensimetrics S14 insert earphones with standard foam tips. Custom filters were applied to flatten the frequency response of the earphones. As in our earlier study^31^, the stimuli were presented with a “Morse code”-like rhythm to enhance their perceptual salience over the sound of the fMRI pulse sequence. Within each trial, 16 short (50-ms) tones and 16 long (200-ms) tones were presented, with each tone having 20-ms raised-cosine onset and offset ramps. Every 700 ms consisted of two short and two long tones, presented in random order, separated by 50-ms gaps. This process was repeated 8 times, with random shuffling of the long and short tones for each repetition, adding up to a total trial length of 5.6 seconds. All tones presented within a single trial had an identical F0 and Fc. Following one trial, after a silent gap with a random duration of uniform distribution between 200 and 600 ms, the next trial was presented.

Within one run of ∼6 minutes, each of the 25 conditions was presented twice, and the 50 trials were presented in a pseudorandomized order, constrained to prevent two identical stimuli from being presented consecutively. Silent periods of 10 seconds were inserted at the beginning of the run and after every 10 trials. A 15-s silence was further appended at the end of the run. These silent periods were added to establish a baseline hemodynamic response^55,56^.

Each participant underwent two experimental sessions of 12 runs each, for a total of 24 runs. Therefore, each condition was presented 24 times per session and 48 times per participant. Multiple repetitions are important for averaging out unwanted measurement variability, and for cross-validation methods of data denoising^53^. Participants were asked to press a button whenever a new trial began to help encourage and monitor the alertness of the participants throughout the experiment. After each run, participants were informed of their behavioral performance (percent correct) in that run. Behavioral performance for all participants is reported in Table S3.

Before the two experimental sessions, participants performed a loudness-adjustment task in a separate scan session, aiming to minimize any loudness differences between the stimuli when presented in the presence of scanner noise. In this task, participants listened to sequences composed of each of the 25 experimental stimuli with the order randomly shuffled, while the scanner was running with the fMRI pulse sequence used in the experimental sessions. Within each sequence, the same stimulus (100 ms) with a silent gap (50 ms) was repeated 20 times with the sound level gradually increasing in 1 dB steps between 43 and 83 dB SPL. Participants were asked to adjust the loudness of the tones to be clearly and comfortably audible over the scanner noise by pressing “1” to decrease the sound level of the sequence and “2” to increase it. Once they were satisfied with the sound level, they could press “3” and the level would remain the same for the rest of the sequence. The task was performed three times for each stimulus. For each participant, the median level of a given stimulus was calculated and later used in that participant’s experimental sessions. In addition to the preliminary loudness adjustments, participants were also instructed to listen for any remaining loudness differences between tones at the beginning of each experimental session. Between the runs, if the participant indicated a need for further adjustments to the sound level, the experimenter would change the sound level slightly (1-3 dB) in the next run based on the participant’s feedback.

##### MRI data acquisition

Both anatomical and functional data were acquired using a Siemens 3T Prisma scanner with a 32-channel head coil at the Center for Magnetic Resonance Research (CMRR) at the University of Minnesota. Functional data were acquired in two experimental sessions on separate days using a continuous gradient-echo EPI sequence (TR = 1800 ms; TE = 30 ms; flip angle = 70°; 2-mm isotropic voxels; number of slices = 60; multiband factor = 3; phase partial Fourier = 6/8). Slices were oriented to be approximately 45° to the brainstem to cover most of the brain and to avoid coverage of the eyeballs whose movement could create Nyquist artifacts within auditory regions. A total of 208 volumes were obtained in each run. Four fieldmaps were collected throughout each experimental session for distortion correction (same slab orientation as the gradient-echo EPI; TR = 400 ms; first TE = 2.16 ms; second TE = 4.62 ms; 4-mm slices).

Anatomical data were acquired in a different session before the experimental sessions. Three T1-weighted and two T2-weighted volumes were collected. MPRAGE T1-weighted parameters included: TR = 2400 ms; inversion time (TI) = 1000 ms; TE = 2.22 ms; flip angle = 8°; 0.8-mm isotropic voxels. T2-weighted parameters included: TR = 3200 ms; TE = 563 ms; 0.8-mm isotropic voxels.

##### MRI data preprocessing

Anatomical and functional data were preprocessed through a series of custom scripts^57^. For anatomical data, gradient unwarping was performed for T1-and T2-weighted volumes to correct for gradient nonlinearity using the gradient coefficient file obtained from the scanner. Then, three T1-weighted volumes were aligned (rigid-body transformation with six degrees of freedom and cubic interpolation) and averaged. The two T2-weighted volumes were aligned in the same way. The averaged T1 was then used for brain segmentation and cortical reconstruction via Freesurfer^28^. The averaged T2 was co-registered to the averaged T1 using rigid-body transformation and cubic interpolation.

Functional data were temporally upsampled from 1.8s to 1s. Fieldmaps were spatially upsampled to a slice thickness of 2mm to match that of the EPI and later used for distortion correction. Distortion correction was performed along with motion correction in a single cubic interpolation. The corrected functional volumes were then co-registered to the averaged anatomical T2 using an affine transformation. Finally, the functional data were re-preprocessed such that the data were resampled onto the mid-grey cortical surface using a single cubic interpolation. The analyses in this paper are performed in surface space, instead of volume space, for better visualization of the cortical mappings of F0 and Fc.

The surface-based functional data were analyzed with GLMsingle^29^, a toolbox for improving the signal-to-noise ratio of estimates of BOLD response amplitudes, especially for experimental designs with a short trial duration and fast-alternating conditions. GLMsingle provides single-trial BOLD response amplitude estimates (“betas”). Upon visual inspection of response timecourses (derived from an independent analysis using a finite impulse response model) and beta estimates, we found that the final run from session 1 of P07 had very weak response amplitudes compared with the rest of the session, and that the response timecourses looked very noisy. Due to the poor data quality, we discarded the data from this run.

##### ROI definition

One ROI was defined for each hemisphere on the participant’s native brain surface space, referred to as the auditory cortex in the results. Aiming to encapsulate the region responsive to the stimuli in the auditory cortex, we manually defined the ROIs as reasonably sized polygons that encompass the majority of voxels with high variance explained by all sound stimuli against the baseline (no stimuli) (see Fig. 2C) or high mean beta value across all conditions around anatomical locations of HG and HS. Importantly, the variance explained by all stimuli against baseline and the mean beta value were computed based on all experimental trials in the session, reflecting the overall brain response towards all sound stimuli played in the experiment (against the baseline). Thus, the ROIs were not biased towards any measure specific to pitch or brightness.

Participant-level ROIs were defined as the union of ROIs defined in the two scanning sessions, respectively. Activation across scanning sessions within individual participants were highly consistent, as demonstrated in Fig. S13. We also defined a group-level ROI for each hemisphere as the intersection of at least 50% of the individual polygons defined above (after mapping each subject to the fsaverage atlas space).

In the ROI-level analyses, aggregating across individual voxels (e.g., Fig. 3B and Fig. 6), we further restricted the ROI to sound-responsive voxels, defined as having a mean beta value significantly larger than 0 (*p* < 0.01).

##### Correction for the effect of sound intensity

Functional data were analyzed first for each individual session. For each voxel, the beta values for all 24 repetitions (for session 1 of P07 whose final run was discarded, 22 repetitions) of a given condition were averaged, forming a tuning surface of 25 data points in a three-dimensional space, with the height of the surface denoting the beta value of each condition, and the x-and y-axes denoting F0 and Fc of the stimuli, respectively. It was observed that the tuning surface averaged across ROIs showed a bias towards tones with low-frequency spectral content (i.e., tones that have low Fc values tend to have stronger BOLD responses), likely resulting from higher sound intensity assigned to low Fc conditions.

Therefore, we decided to correct for such a bias based on the intensity-beta linear fit for each participant (Fig. S14). First, voxels that were both within the ROI polygon and sound-responsive (see *ROI definition*) were selected as “good voxels”. Second, the mean betas across all repetitions for each of the 25 conditions were averaged across these “good voxels” for each session. Third, the 50 intensity-beta pairs from the two functional sessions of one participant were pooled to obtain a linear fit between sound intensity and mean beta from “good voxels” for each participant. Finally, the intensity-corrected beta estimate of every single trial in each of the two sessions was calculated by dividing the original beta of this trial by the predicted beta from the intensity-beta linear fit of this participant at the sound intensity of this trial.

##### Fitting and evaluating encoding models

The following five Gaussian-based encoding models were designed to characterize the tuning of each voxel to F0 and Fc after the two dimensions were logarithmically transformed: 1) The *interaction* model assumes that each voxel is tuned to both F0 and Fc, and the preferred frequency of one dimension is dependent on the value of the other dimension; 2) the *independent* model assumes that each voxel is tuned to both F0 and Fc and that the two dimensions are processed independently (i.e., the preferred F0 of the voxel does not depend on the stimulus Fc); 3) the *Fc-tuned* model assumes that the voxel is tuned only to Fc and is invariant to F0; 4) the *F0-tuned* model assumes that the voxel is tuned only to F0 and is invariant to Fc; and 5) the *untuned* model assumes that the response of the voxel is invariant to both F0 and Fc. **Error! Reference source not found.** provides the equations for each of the models. In models 1-4, *f*_Fc_ and *f*_F0_ represent the stimulus F0 and Fc values, CF_F0_ and CF_Fc_ represent the coordinates of the peak of the tuning surface, and s represents the width (standard deviation) of the Gaussian curve in a given dimension. *g* represents the height of the tuning surface. Among the five models, the comparison between the *independent* and the *interaction* models is of most interest for this study, as this comparison tests the hypothesis that tuning in one dimension is affected by the other dimension. Mathematically, these two models are highly similar, in that the tuning surface of the *interaction* model is just that of the *independent* model rotated by θ degrees on the Fc-F0 plane (see schematic illustration in Fig. 3A). We note that the value of θ itself is complicated to interpret. This is because our 5×5 stimulus grid does not necessarily span the full range of voxel’s response variation (from low response to high response and to low response again). With only partial sampling, different models fits with different θ values may fit equally well.

Before model fitting, the intensity-corrected beta estimates from two sessions in one participant were combined, forming a participant-level dataset. This was done by averaging the intensity-corrected beta estimates from session 1 and session 2 on the trial level so that the participant-level dataset had the same structure (i.e., number of repetitions for each condition, number of runs, and number of trials) as one single session. To balance the amount of data fed into the participant-level dataset between the two sessions, the last run of session 2 for P07 was discarded, along with session 1 for P07 which had data quality issues. As a result, each voxel had 12 (11 for P07) tuning surfaces, each from a quasi-run in the participant-level dataset consisting of 25 beta estimates for 25 conditions. The tuning surfaces were then averaged across quasi-runs. This averaged tuning surface was then used to fit the five encoding models.

For each voxel, the performance of each of the five encoding models was evaluated using an N-fold leave-one-run-out cross-validation procedure (N is the number of runs in the current participant-level dataset). First, averaged data across N-1 runs were used to find the training parameters. The parameter *g* was imposed to be non-negative. Other parameters were not constrained. To avoid local minima, the parameters were initialized with various sets of initial seeds based on the voxel response. MATLAB’s lsqcurvefit function was used to optimize for least-square solutions. The fitted parameters were then tested on the held-out run to obtain an R^2^ value (coefficient of determination, calculating by subtracting the proportion of the original data variability that is not explained by the model fit from 1). This procedure was repeated N times, each holding out a different run as the testing set, yielding N R^2^ values. The average of these N values was then calculated as the cross-validation R^2^ for this model. Higher R2 indicates better model performance.

#### Behavioral Experiment

##### Participants

All ten participants from the MRI experiment took part in the behavioral experiment, along with 10 newly recruited participants [8 females, 2 males, mean (SD) age of 24.8 (4.2) ranging from 19 to 30 years] from the University of Minnesota community. All of the newly recruited participants had normal hearing according to the criteria in the MRI experiment. Three participants were identified as non-musicians, having two or less years of formal musical training. The other seven participants had an average of 11.8 years of musical training on non-percussion instruments. Six of the ten new participants were tone-language speakers (Mandarin Chinese and/or Cantonese). The experimental protocol was approved by the University of Minnesota Institutional Review Board (Approval No.: 0605S85872). All participants provided written informed consent before the experiment.

##### Stimuli

The stimuli were harmonic complex tones, digitally generated in the same way as in the MRI Experiment, except for the duration of the tones. Each tone in the behavioral experiment had a duration of 200 ms, including 20-ms raised-cosine onset and offset ramps. Each trial contained two tones played sequentially, separated by a silent gap of 300 ms. All the stimuli were converted to analog at a sampling rate of 48 kHz using a Lynx E22 sound card with 24-bit resolution and were presented diotically via open-ear headphones (Sennheiser HD 650) in a sound-attenuating booth at a level of 60 dB SPL.

Before the main experiment, participants’ F0 difference limens (DL_F0_) and Fc difference limens (DL_Fc_) were measured for the middle tone in the 5×5 stimulus set used in the MRI Experiment (F0 = 420 Hz and Fc = 1500 Hz). In the measurement of DL_F0_, Fc was held constant at 1500 Hz while the F0 of the two tones was different and geometrically centered around 420 Hz. In the measurement of DL_Fc_, F0 was held constant at 420 Hz while the Fc of the two tones was different and geometrically centered around 1500 Hz. The individual DL_F0_ and DL_Fc_ from this preliminary experiment were used to set the DLs for each listener in the main experiment.

In the main experiment, as in the DL measurements, all variations in F0 and Fc were geometrically centered around F0 = 420 Hz and Fc = 1500 Hz. For the up-down judgment of pitch, the ΔF0 between the two tones was always set to one DL_F0_ for each listener individually, whereas ΔFc was set to-2,-1, 0, +1, or +2 DL_Fc_, where positive values imply congruence between the F0 and Fc direction, negative values imply incongruence, and 0 implies no change in Fc. For the up-down judgment of timbral brightness, ΔFc between the two tones was always set to one DL_Fc_ for each listener individually, whereas the ΔF0 was set to-2,-1, 0, +1, or +2 DL_F0_.

##### Procedure

Both the preliminary DL measurement and the main experiment adopted a two-interval two-alternative forced-choice paradigm delivered through the AFC software package^58^. Two virtual buttons labeled “1” and “2” were displayed on the computer screen, indicating the first and the second tones, respectively. Each button flashed when the corresponding tone was played. Participants were asked to select which tone was higher in pitch or brightness (depending on the condition), and responded by clicking on one of the two buttons displayed on the screen with a mouse or by pressing “1” or “2” on the keyboard. After each response, immediate feedback of “Correct” or “Incorrect” was displayed on the screen before proceeding to the next trial. Feedback was provided in both the initial DL measurements and the main experiment.

The DL measurements used a standard two-down one-up tracking procedure that tracks the 70.7% correct point on the psychometric function^59^. Each track began with a DF (DF0 in the measurement of DL_F0_ and DFc in the measurement of DL_Fc_) of 20% (of the lower tone), which decreased or increased by a factor of two, depending on whether the response of the previous trial was correct or not. After the first reversal from an “up” to a “down” direction of the tracking procedure, DF changed by a factor of 1.41. After the second such reversal, DF changed by a factor of 1.19. The tracking stopped after a further four reversals, and the DL of this track was calculated as the geometric mean of the DF values at the last four reversal points.

Before the measurements, participants were given examples of tones with low and high pitches, as well as tones with duller or brighter timbres. They were also encouraged to use the immediate feedback given in the experiment to establish their concepts of pitch and timbral brightness. Each participant completed four measurements of DL_F0_ and four measurements of DL_Fc_ in a pseudorandomized order in about 10 minutes. Each participant’s final DL_F0_ and DL_Fc_ were calculated as the geometric mean of the four measurements.

The main experiment included 10 blocks of the up-down judgments of pitch and 10 blocks of the up-down judgments of timbral brightness in a pseudorandomized order. As in the DL measurements, participants were asked to indicate which of the two tones had a higher pitch (for the blocks of pitch judgment) or brighter timbre (for the blocks of brightness judgment). Each block had five ΔFs (-2,-1, 0, 1, and 2 DLs, linearly spaced) in the non-target dimension (DFc in pitch judgments and DF0 in brightness judgments) each with 10 trials presented in a random order, and the DF in the target dimension (DF0 in pitch judgment and DFc in brightness judgment) was held constant at the participant’s DL in this dimension.

There were also 10 practice trials at the beginning of each block, where the DF in the target dimension was set to 2 DLs instead of one, to help participants find the perceptual cues needed to succeed in the task. Therefore, each block contained a total of 60 trials (two practice trials and 10 test trials at each value of DF in the non-target dimension). Each participant took about 60 minutes to complete the main experiment.

##### Data and statistical analysis

As described in *Stimuli*, the geometric means of the DL measurements were used as the DL_F0_ and DL_Fc_ in the main experiment. For the main experiment, percent correct for each of the five conditions (ΔF_non-target dimension_ =-2,-1, 0, 1, or 2 DLs) in the two tasks (pitch and brightness direction discrimination) was each calculated and converted to *d’* using the equation

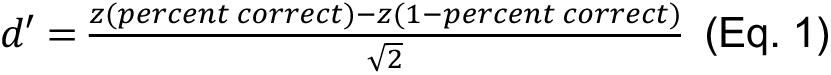

for a 2AFC task^60^. The sensitivity index *d’* was used instead of percent correct because *d’* is proportional to ΔF in linear units^61^. A linear function was then fitted to the five *d’* values from each the two tasks, with the x-axis being ΔF_non-target dimension_ and the y-axis being the *d’* in this condition. The slope of the function therefore reflected the effect of the variation in the non-target dimension on the discriminability of the target dimension. To evaluate the contribution of the non-target dimension to discrimination of the target dimension, the slope of the linear fit was divided by the theoretical *d’* that corresponds to one DL, i.e., *d’* of 1.09 for an accuracy of 70.7%^60^. Two-tailed Wilcoxon signed-rank tests were performed to compare the weights of the non-target dimension with a zero-slope null hypothesis (i.e., no influence of the non-target dimension). Group averaged *d’* values and their linear fits were also calculated and are shown in Fig. 1A.

