## Supplemental Materials for "Cortical substrates of perceptual confusion between pitch and timbre"

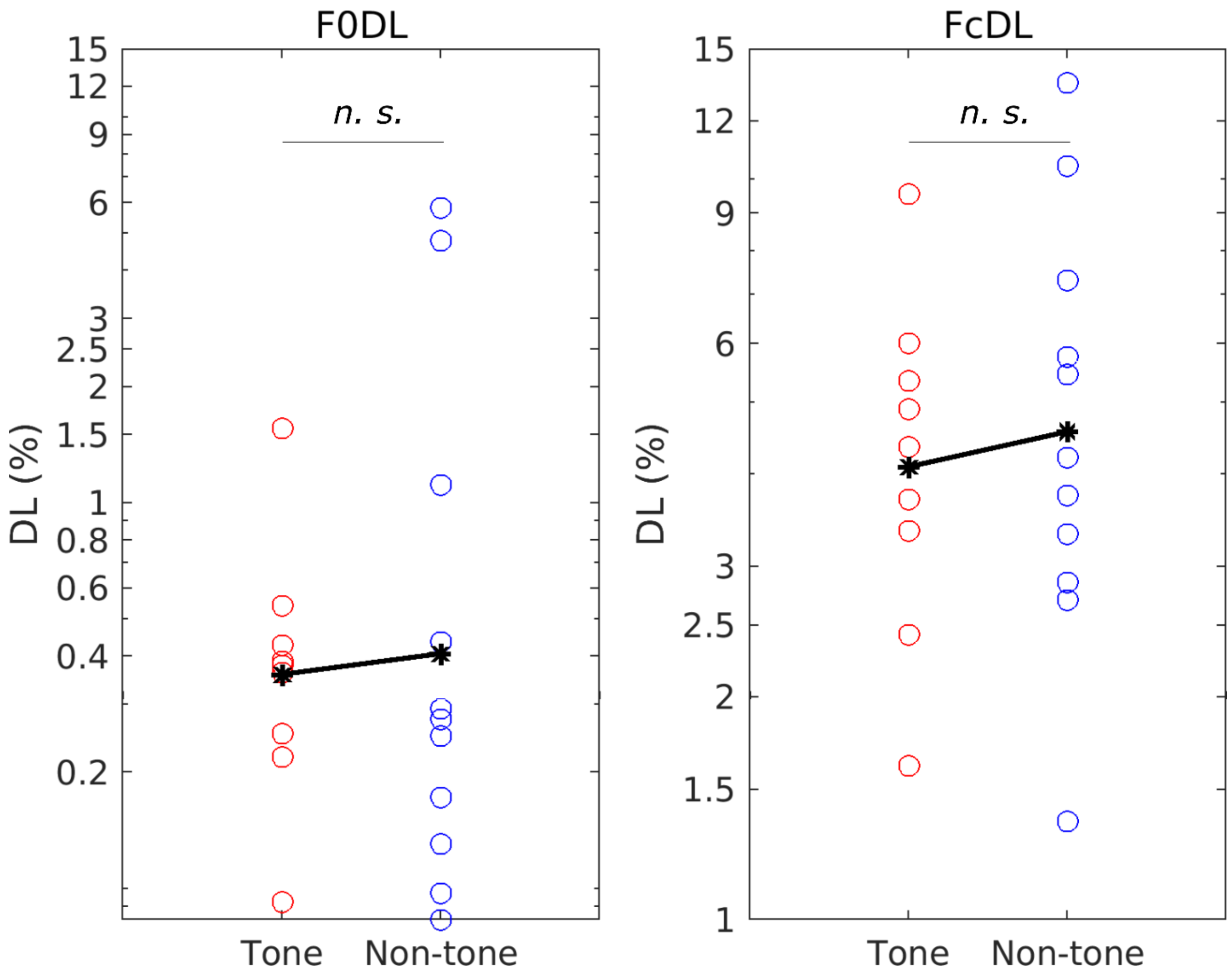


**Fig. S1. No difference in either $DL_{F0}$ or $DL_{Fc}$ between tone and non-tone language speakers.** Mann-Whitney U test: $N_{tone}$ = 9, $N_{nontone}$ = 11; $U_{F0}$ = 100, $p$ = 0.71; $U_{Fc}$ = 90, $p$ = 0.77.

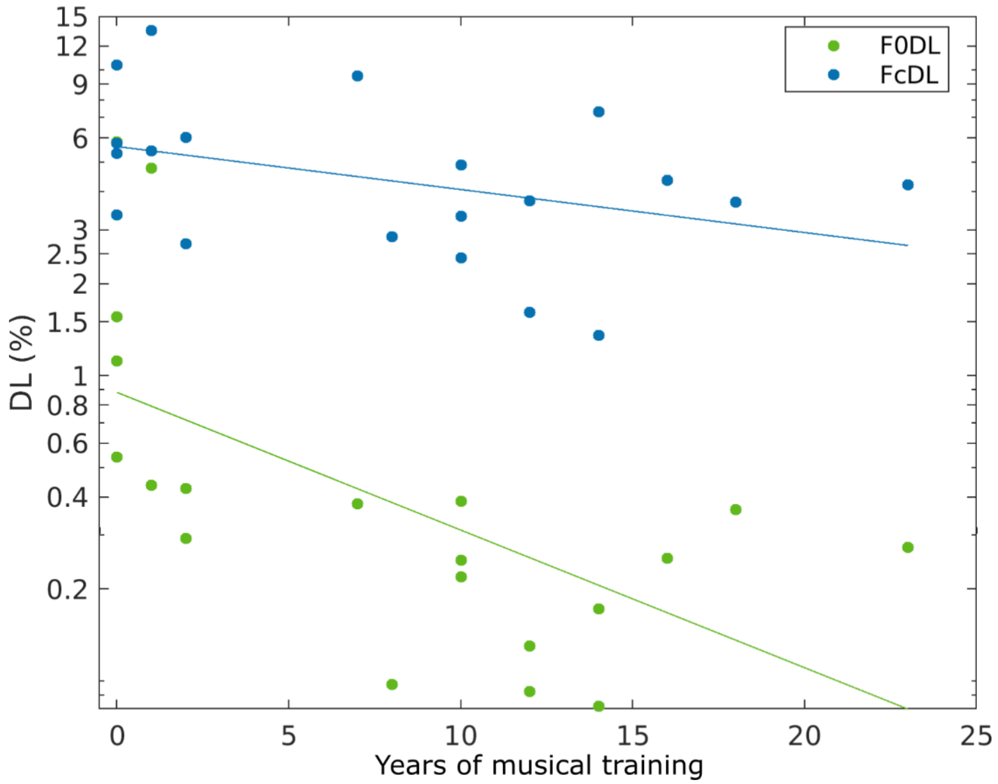


**Fig. S2. Effect of musical training on $DL_{F0}$ and $DL_{Fc}$.** Year of music training was negatively correlated with $DL_{F0}$ but not with $DL_{Fc}$ (Spearman's correlation: $N = 20$; $\rho_{F0} = -0.744$, $p < 0.001$; $\rho_{Fc} = -0.397$, $p = 0.082$).

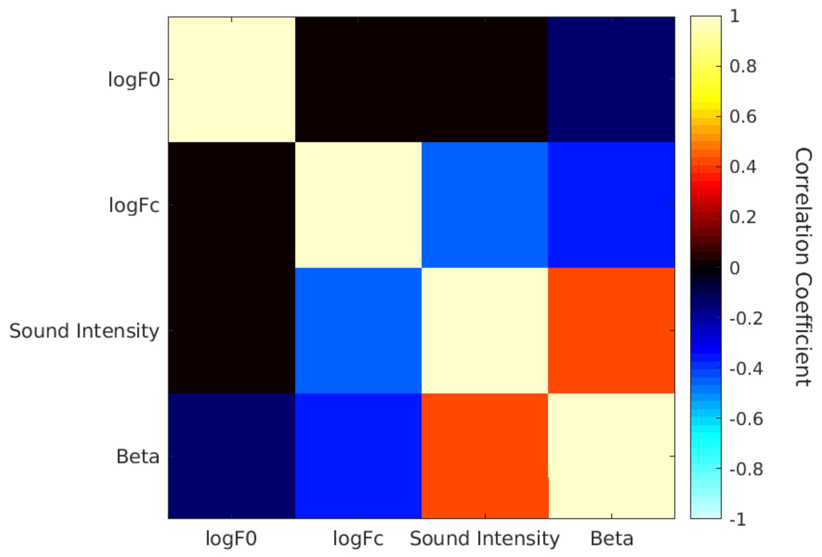


**Fig. S3. BOLD response bias towards low Fc conditions introduced by correlation with sound intensity.** Correlation matrix between sound intensity assigned to each of the 25 conditions in each experimental session, their corresponding F0 and Fc values in logarithmic scale ("logF0" and "logFc"), and their corresponding BOLD response averaged across all "good voxels" ("Beta") using Pearson correlation. There is a strong positive correlation between sound intensity and BOLD response. High sound intensity tended to be assigned to conditions with low Fc values. These correlations led to a strong negative correlation between Fc values and BOLD responses, introducing a BOLD response bias towards low Fc, which distorted the neural tuning pattern.

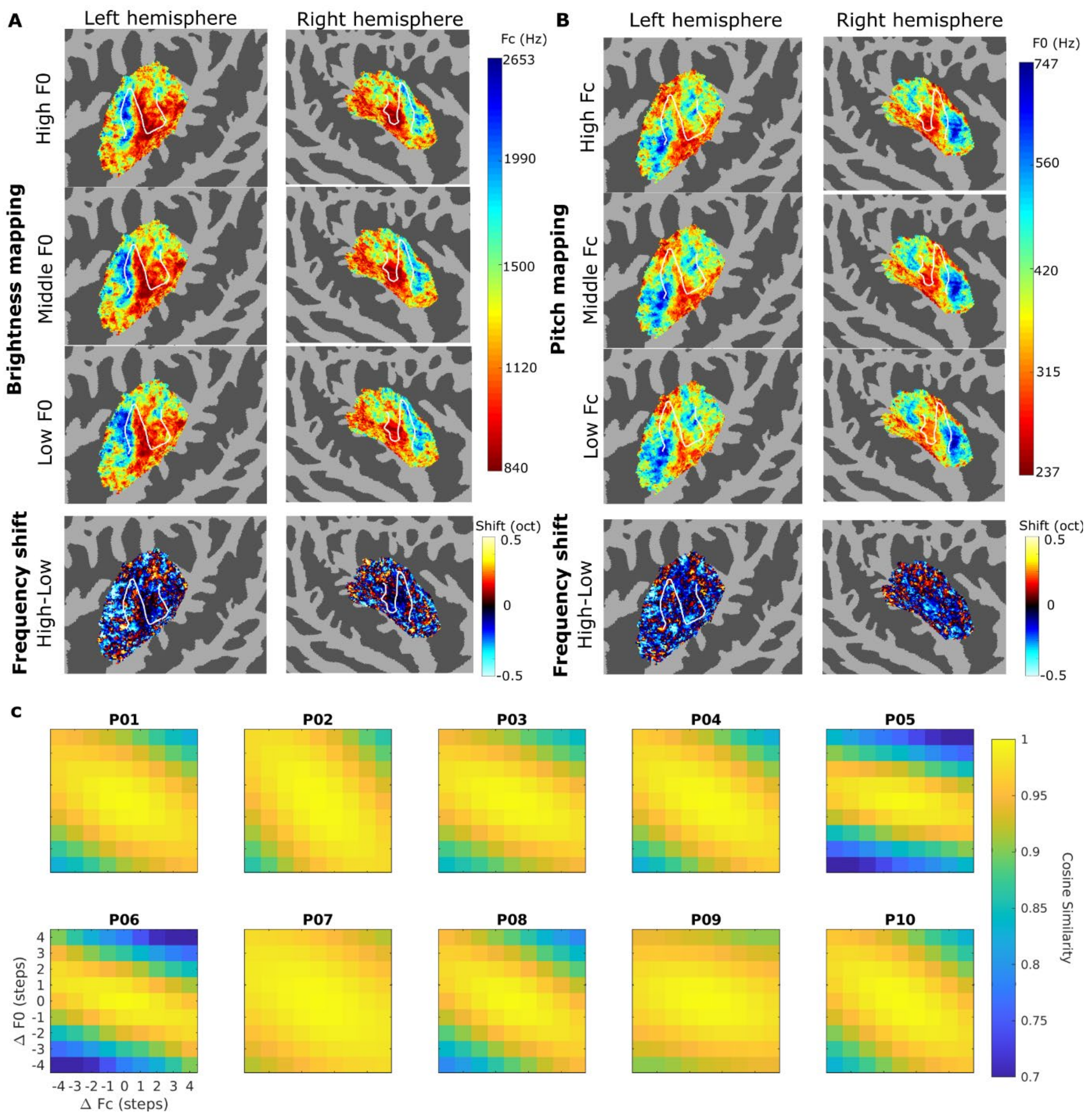


**Fig. S4. Cortical substrates of pitch-brightness confusion are replicated in the data without sound-intensity correction. (A)** Replicating the within-voxel preferred Fc shift in Fig. 4A, showing an overall downward shift in preferred Fc as F0 increases. **(B)** Replicating the within-voxel preferred F0 shift in Fig. 4B, showing an overall downward shift in preferred F0 as Fc increases. **(A)-(B)** Replicating the global similarity of the tonotopic gradients between brightness mapping and pitch mapping in Fig. 5. **(C)** Replicating population-level representational similarity matrix in Fig. 6, showing that congruent changes between F0 and Fc elicited less similar neural activation patterns than incongruent changes.

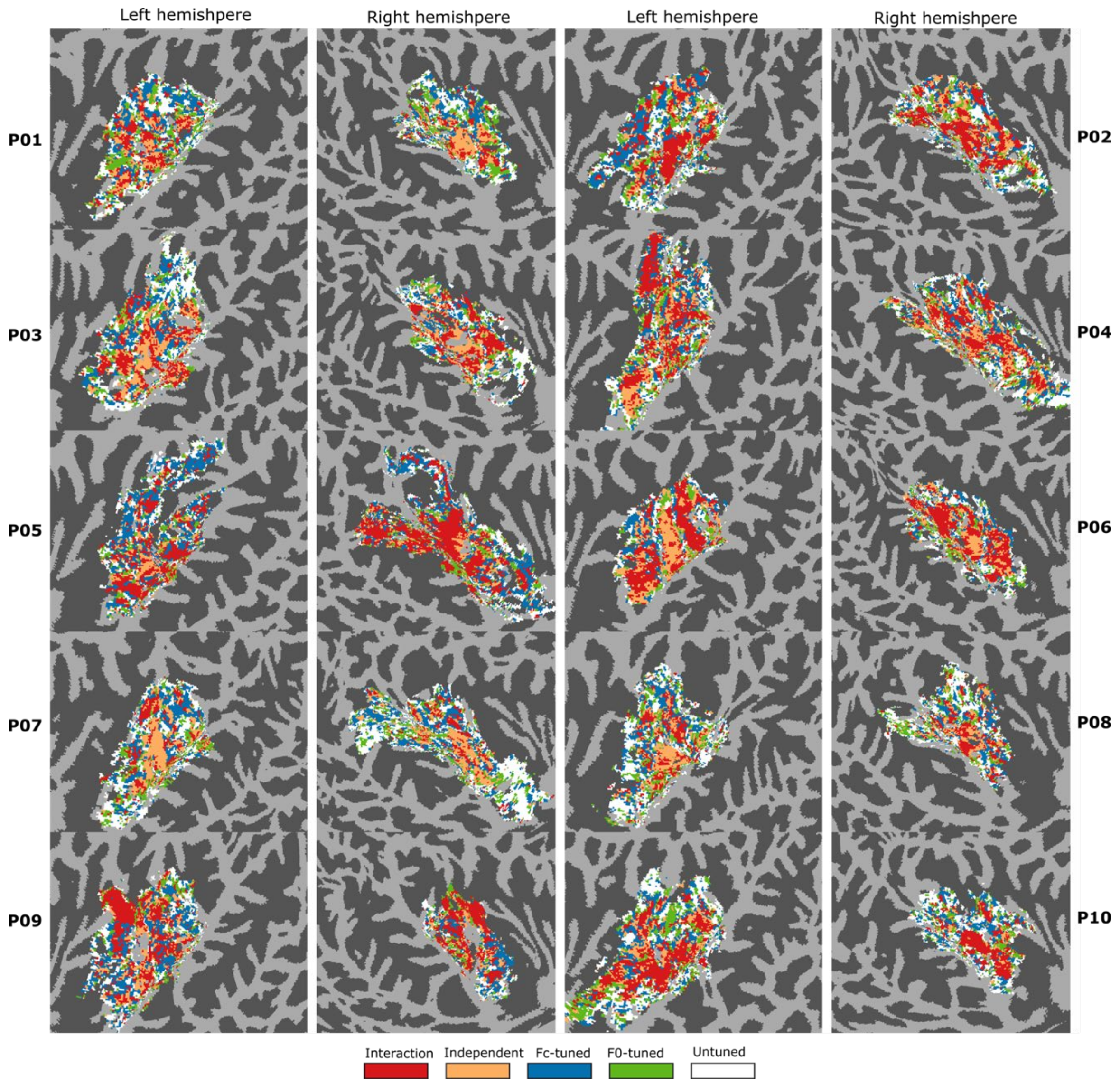


**Fig. S5. Spatial distribution of voxels with different tuning properties ("best model map") for all participants** Individual results of Fig. 3C. Color of the voxels within the ROI indicates the best-performing model for this voxel. The first two columns are left and right hemispheres, respectively, of Participants 1, 3, 5, 7, and 9. The last two columns are left and right hemispheres, respectively, of Participants 2, 4, 6, 8, and 10. Only voxels within the ROI with a mean beta value significantly larger than 0 ($p < 0.01$) are shown (same for Fig. S6 and Fig. S11).

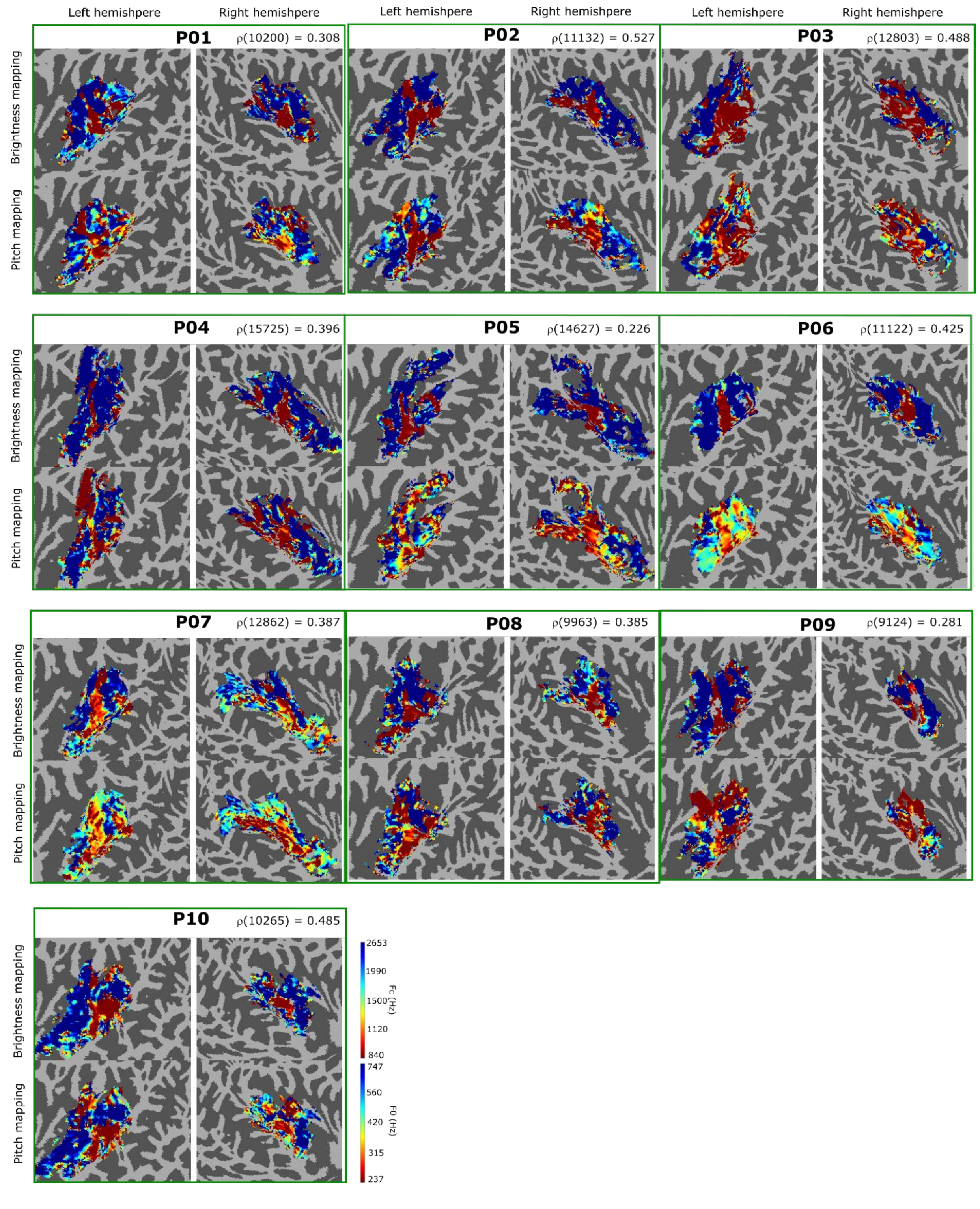


**Fig. S6. Brightness and pitch mappings of all participants** Individual results of Fig. 5. Each green box encloses one participant's brightness mapping (top row) and pitch mapping (bottom row), showing both left hemisphere (left column) and right hemisphere (right column). Top row: preferred Fc at the middle F0 (420 Hz). Bottom row: preferred F0 at the middle Fc (1500 Hz). ρ: Spearman's correlation coefficient. *P*s < 0.001 for all ten participants.

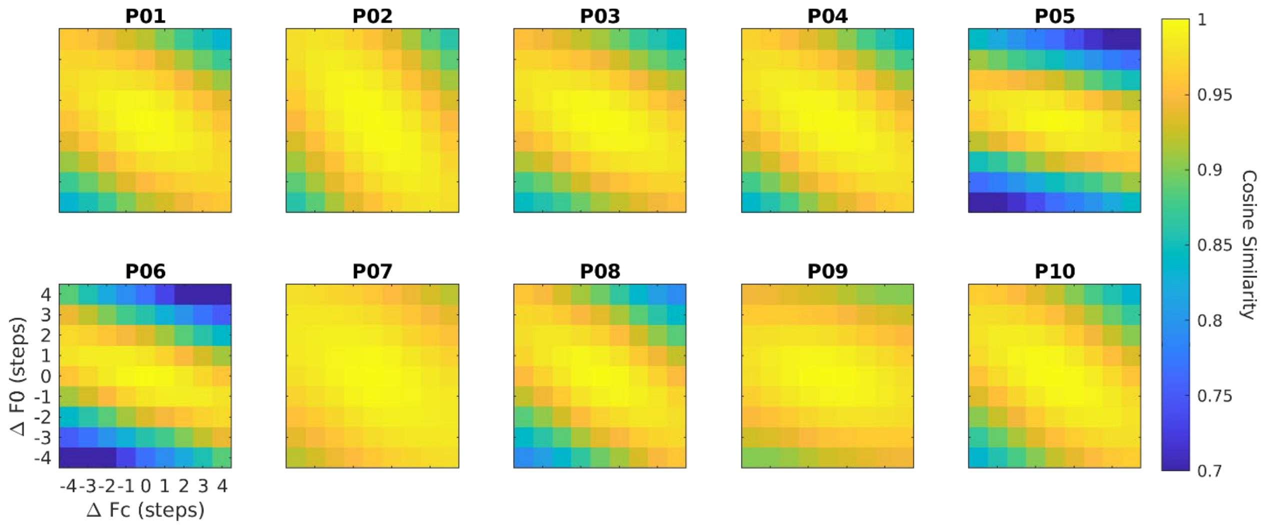


**Fig. S7. Similarity matrices showing cosine similarity between pairs of conditions for all participants.** Individual results of Fig. 6. Description of the computation of the similarity matrix can be found in the main manuscript.

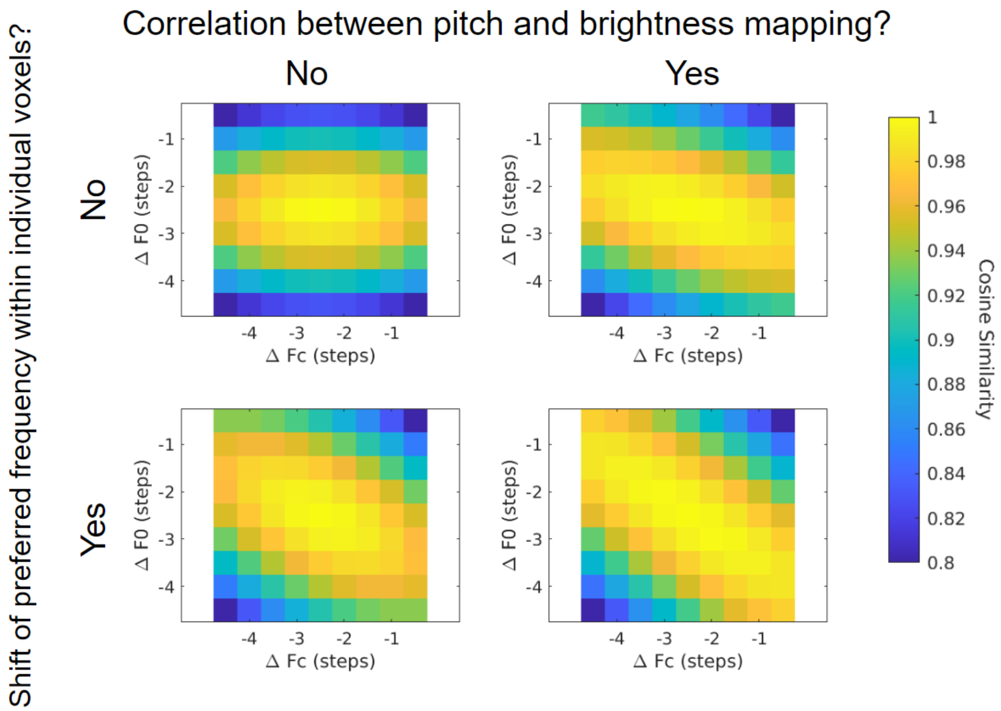


**Fig. S8. Both correlation between pitch and brightness mappings and preferred frequency shift within individual voxels can contribute to the tilt of the similarity matrix** A simulation was performed to investigate the contribution of the two distinct lines of evidence: shift of preferred frequency within individual voxels (see Within-voxel substrates of perceptual confusion) and correlation between pitch and brightness mappings across voxels (see Global cortical substrates of perceptual confusion) to the Combining within- and across-voxel patterns to predict perceptual confusion. Preferred Fc's and preferred F0s of all voxels within the group-level ROIs were used as the parameters $CF_{F0}$ and $CF_{Fc}$ along with parameters $\sigma_x = 1$, $\sigma_y = 1.5$, and $g = 3$ (see Model No. 1 in Table 1) to generate model fits for these voxels. The combination of parameters was chosen to represent the typical tuning patterns of "good voxels". To test the contribution of the evidence in the within-voxel tuning, model parameter $\theta$ was set to 0 (no shift in preferred frequencies, top row) or 30 (shift in preferred frequency, bottom row). To test the contribution of the evidence in the across-voxel mappings, the preferred Fc values were shuffled across voxels so that they were no longer correlated with the preferred F0s (left column). Model fits in the four conditions were then used to calculate the similarity matrix. As the figure shows, the similarity matrix did not show a downward tilt when neither of the factors were present (top left). Both factors can create the tilt (top right and bottom left) and their effects were additive (bottom right).

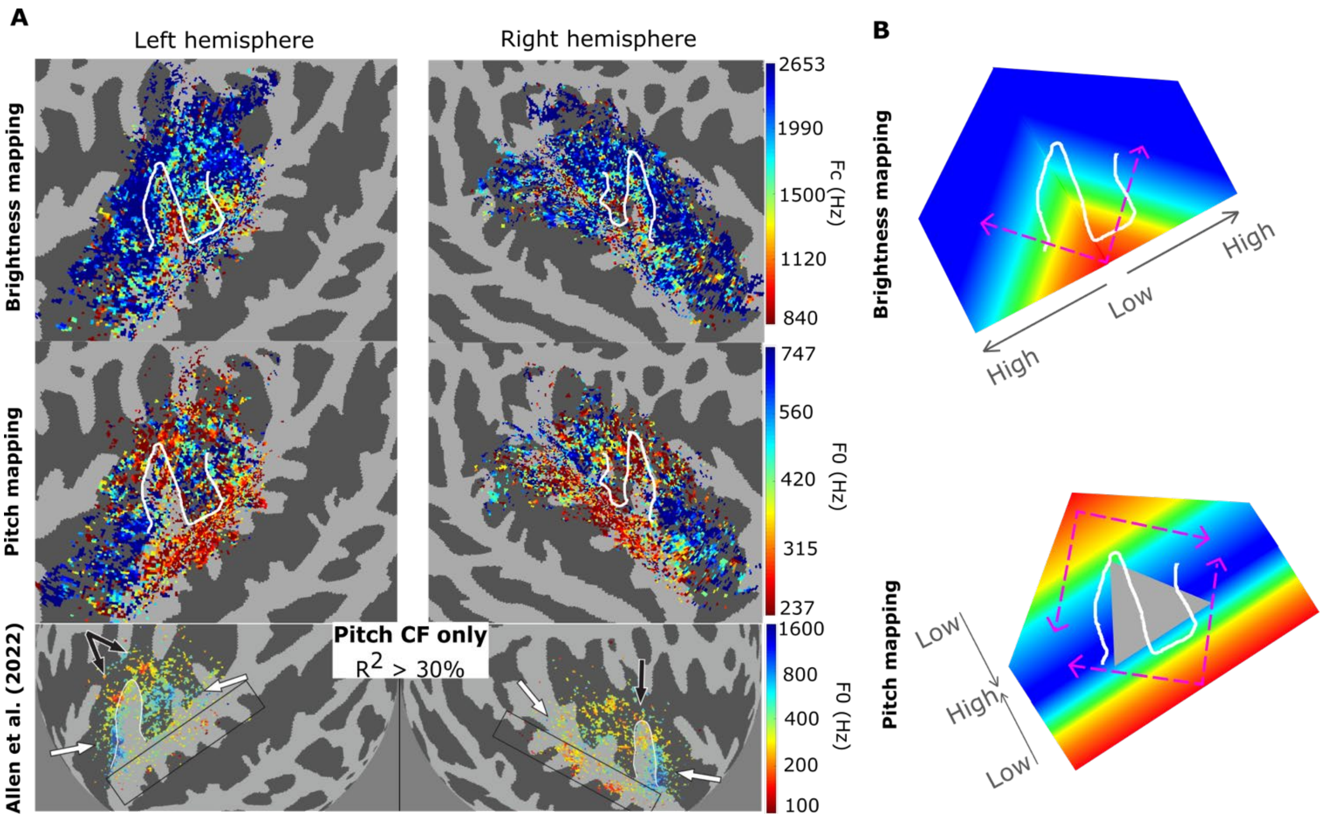


**Fig. S9. Tonotopic patterns in the exclusive tuning to brightness and pitch. (A)** Top row: Preferred Fc at the middle F0 (420 Hz) for voxels that were exclusively tuned to Fc but not to F0, pooled across all participants. Middle row: Preferred F0 at the middle Fc (1500 Hz) for voxels that were exclusively tuned to F0 but not to Fc, pooled across all participants. Bottom row: Preferred F0 for voxels that were tuned to F0 at an Fc of 2400 Hz pooled across all participants, data and figure from [S1]. Note that top and middle rows are flattened surface maps, whereas the bottom row shows the cortical surface in a spherical space; also, the color ranges were set differently according to the frequency range used in these studies in order to better visualize the gradient transitions. These differences in visualization should be taken into account when comparing within and across studies. **(B)** Schematic depiction of the tonotopic gradients across the auditory cortex observed from this study and [S1]. Grey and magenta arrows point from low-frequency-preferred regions to high-frequency-preferred regions. The grey triangle indicates the region that is not exclusively tuned to pitch. White contours delineate the boundaries of HG and HS. Only the left hemisphere is shown.

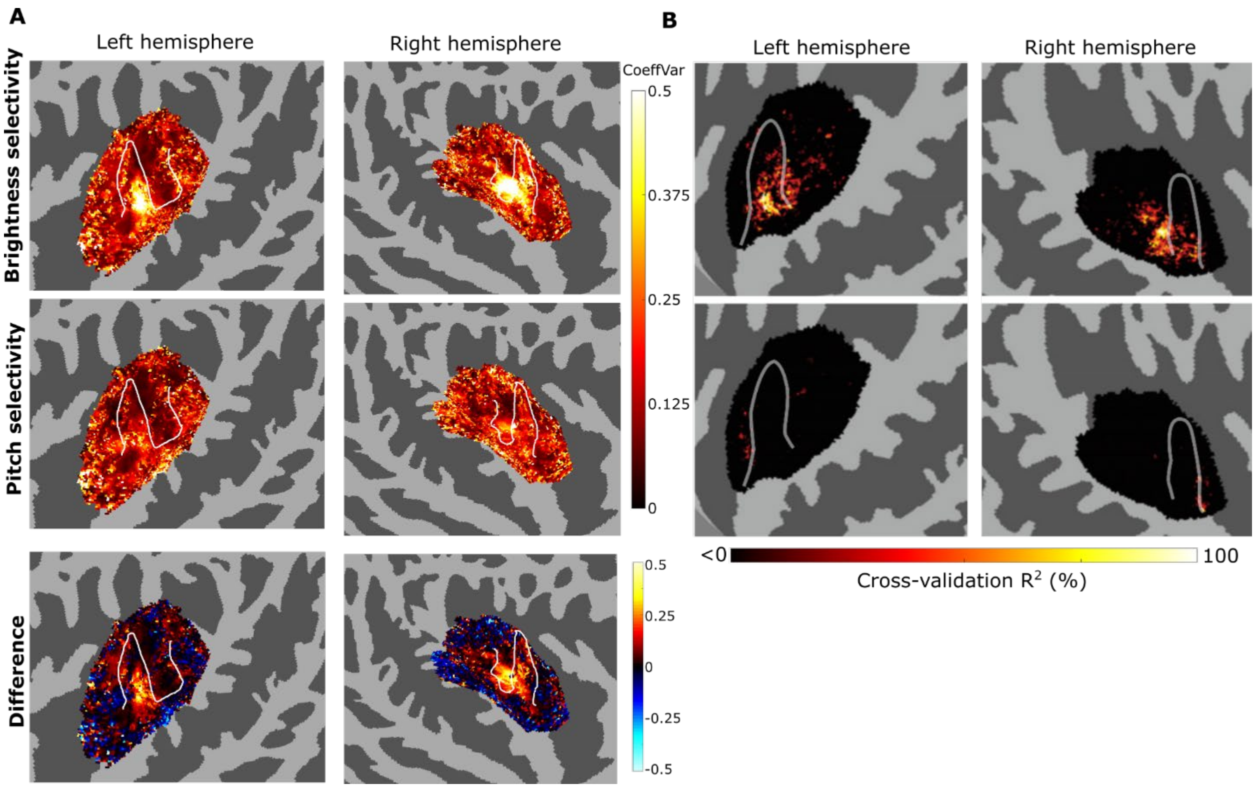


**Fig. S10. Cortical selectivity to brightness is stronger than to pitch in the core regions but somewhat weaker in the surrounding region. (A)** Selectivity to pitch and brightness observed in this study. Top row: selectivity to brightness, quantified by the standard deviation of the Fc tuning curve averaged across F0 divided by the mean of the curve. Middle row: selectivity to pitch, quantified by the standard deviation of the F0 tuning curve averaged across Fc divided by the mean of the curve. Bottom row: brightness selectivity subtracted by pitch selectivity. CoeffVar: coefficient of variation. **(B)** Selectivity to brightness (top row) and to pitch (bottom row) observed in [S1], quantified by the cross-validation $R^2$ of the fitted Gaussian-shaped Fc tuning curves at an F0 of 200 Hz and that of F0 tuning curves at an Fc of 2400 Hz. Figure adapted from [S1]. Black and white contours delineate the boundaries of HG and HS.

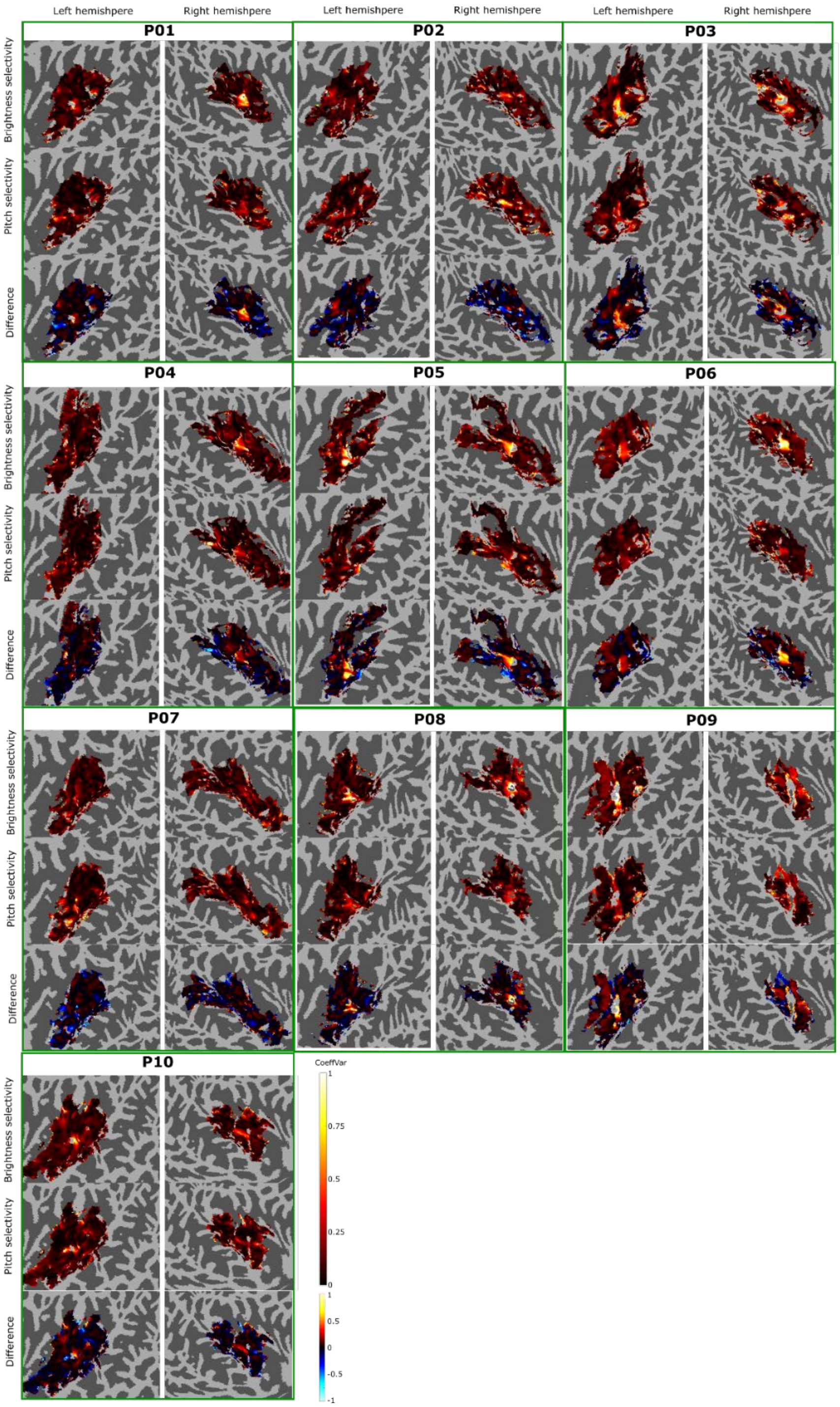


**Fig. S11. Cortical selectivity to brightness and pitch for all participants** Individual results of Fig. S10. Each green box encloses one participant's selectivity to brightness (top row) and selectivity to pitch (bottom row), showing both left hemisphere (left column) and right hemisphere (right column). Top row: selectivity to brightness, quantified by the standard deviation of the Fc tuning curve averaged across F0 divided by the mean of the curve. Middle row: selectivity to pitch, quantified by the standard deviation of the F0 tuning curve averaged across Fc divided by the mean of the curve. Bottom row: brightness selectivity subtracted by pitch selectivity.

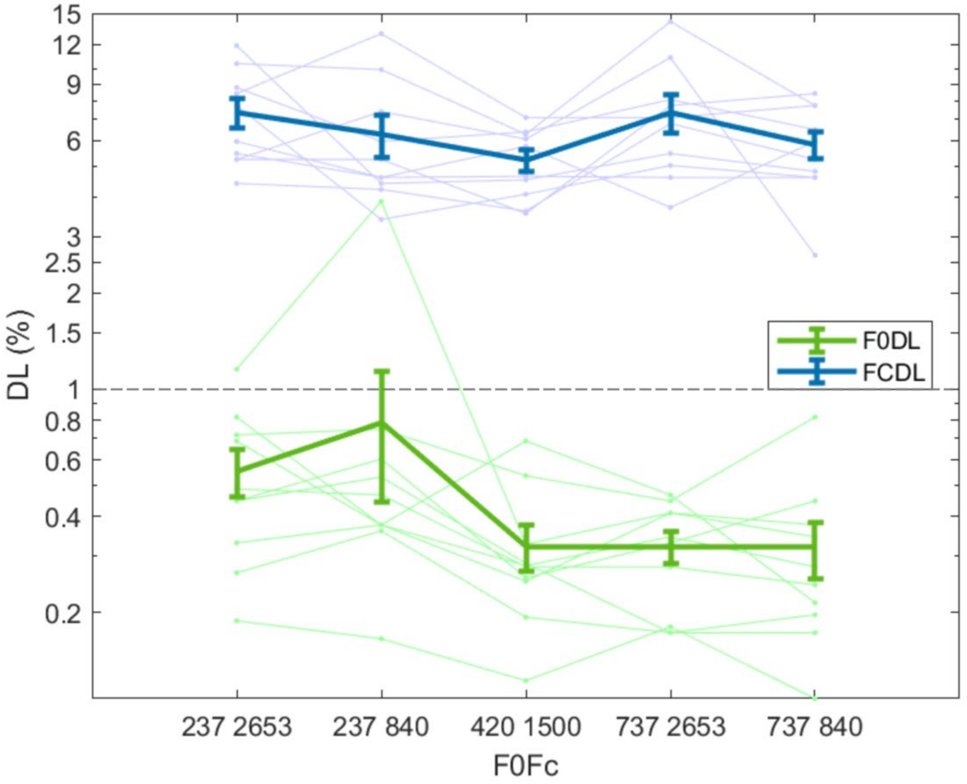


**Fig. S12. Difference limens were comparable across experimental conditions**. Line plots show $DL_{F0}$ (green lines) and $DL_{Fc}$ (blue lines) of a subset of five conditions measured from participants in the MRI experiment. The four corners and the center of the stimulus grid (see Fig. 2B) were chosen to represent different theoretical levels of resolvability across the conditions, with the leftmost condition being the least resolved and the rightmost condition the most resolved. Light-colored circles are data from individual participants, connected with light-colored lines (light green: $DL_{F0}$ ; light blue: $DL_{Fc}$ ). Colored lines are group means. Error bars are standard errors across participants. A non-parametric Friedman test indicated a significant effect of condition [$DL_{F0}$: $\chi^2(4) = 15.63$, $p = 0.004$; $DL_{Fc}$: $\chi^2(4) = 10.90$, $p = 0.03$], with only the $DL_{F0}$ of one pair of tones (F0 = 237 Hz, Fc = 2653 Hz and F0 = 420 Hz, Fc = 1500 Hz) showed a significant difference (two-tailed Wilcoxon signed-rank test with Bonferroni correction: N = 10, W = 51, $p = 0.014 > 0.05/10$). Nevertheless, in all cases, the mean $DL_{F0}$ was less than 1%, indicative of accurate pitch perception (and access to spectrally resolved components) in all conditions.

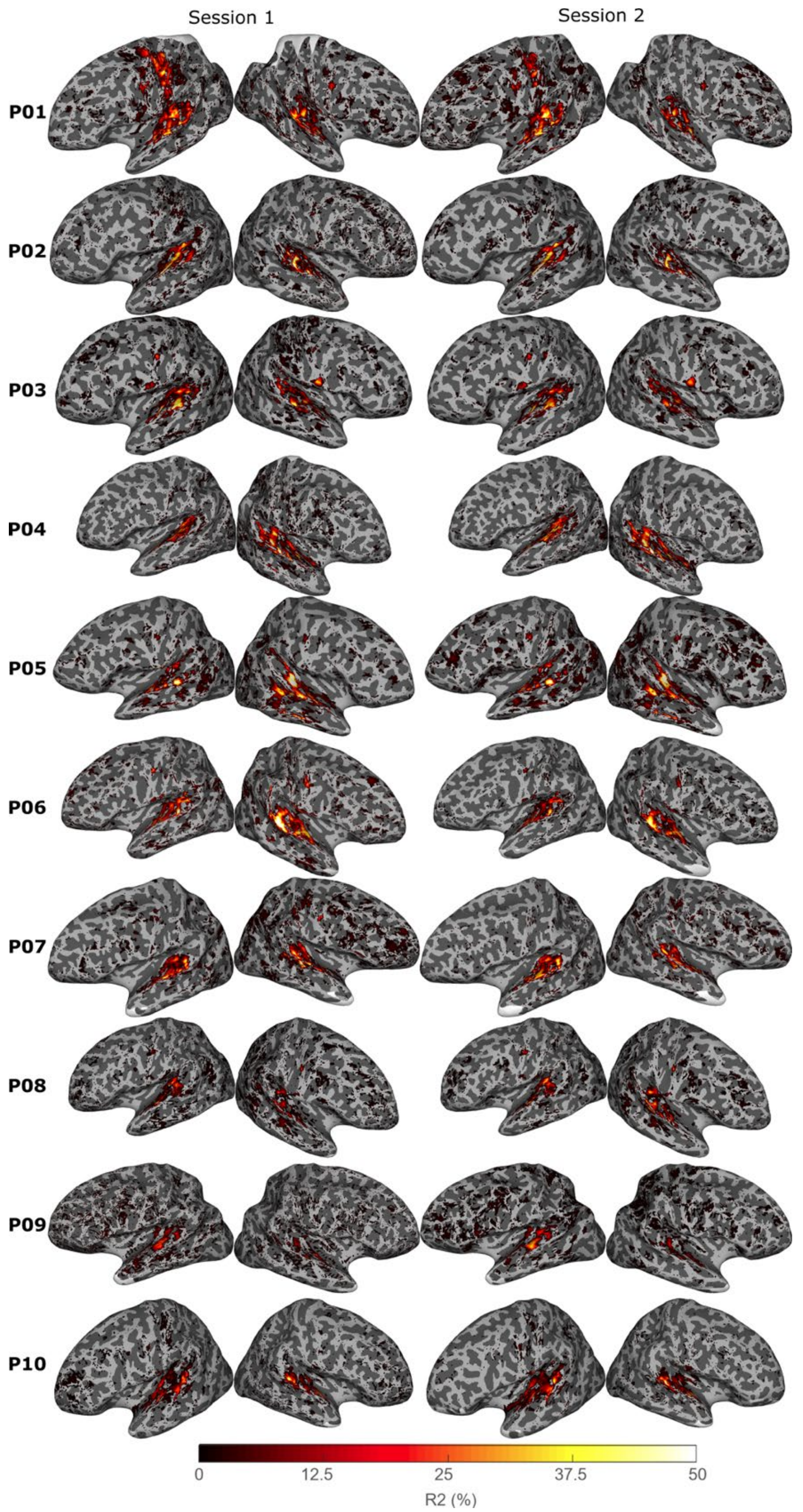


**Fig. S13. Variance explained by all experimental conditions against baseline silence is consistent across scanning sessions.** Each row represents data from the first and second scanning sessions of the same participant. The threshold of each map was computed by fitting two Gaussian Mixture Model with n=2 to the data across the brain in this scanning session to find the point at which the posterior probability is equal across the two Gaussians.

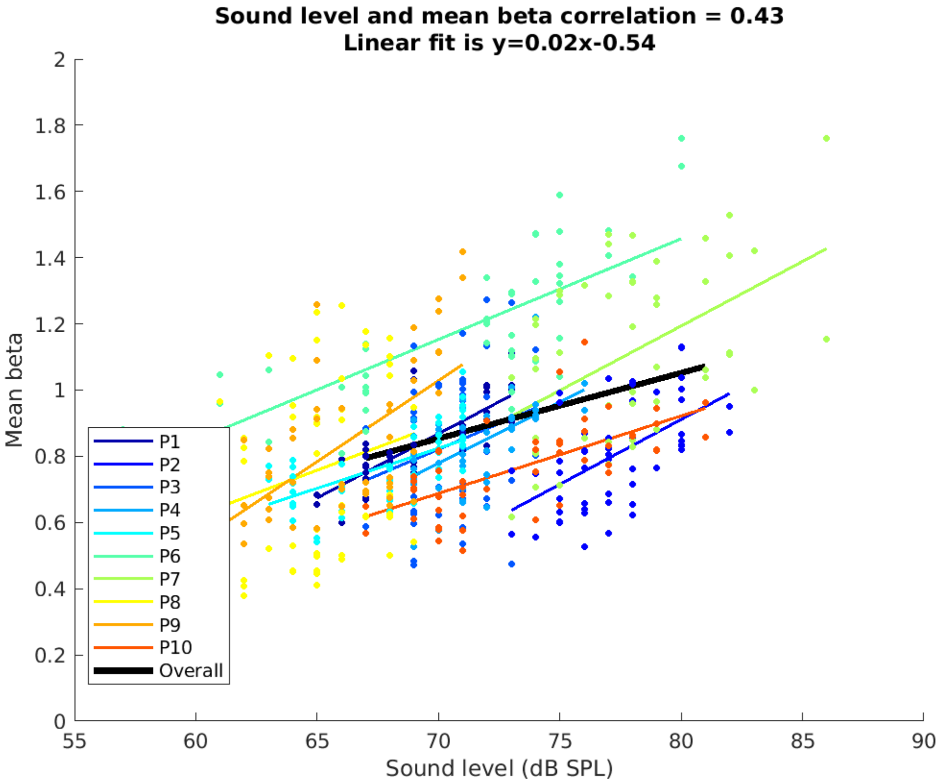


**Fig. S14. Positive effect of sound intensity on cortical activation.** Each dot represents a beta value averaged across all voxels that are both within the ROI polygon and have a mean beta significantly larger than 0 ($p < 0.01$) for one of 25 conditions from one scanning session. All 50 dots from one participant were used to get the linear fit between sound level and mean beta for each participant. Colored lines represent linear fits of individual participants, which were later used to correct for the effect of sound intensity on beta values. The black line represents the linear fit using intensity-beta combinations from all participants (i.e., all dots on the scatter plot).

**Table S1. Number of voxels with a negative or positive shift in preferred Fc and F0.**

| | Preferred Fc | | Preferred F0 | |
|---|---|---|---|---|
| Participant | Negative Shift | Positive Shift | Negative Shift | Positive Shift |
| P01 | 2268 | 2329 | 3152 | 3184 |
| P02 | 3250 | 1406 | 4373 | 1958 |
| P03 | 2766 | 1858 | 4024 | 2755 |
| P04 | 2596 | 2717 | 3314 | 3131 |
| P05 | 5522 | 802 | 8460 | 1272 |
| P06 | 2071 | 2090 | 5196 | 3959 |
| P07 | 5377 | 3501 | 5911 | 3849 |
| P08 | 3439 | 1642 | 3650 | 1782 |
| P09 | 2400 | 866 | 3453 | 1380 |
| P10 | 4751 | 1033 | 5602 | 1193 |

**Table S2. Musical and language backgrounds of individual participants.**

| Participant | Years of Music Training | Tone Language Speaker? | Multilingual? |
|---|---|---|---|
| P01 | 2 | No | Yes |
| P02 | 12 | Yes | Yes |
| P03 | 16 | Yes | Yes |
| P04 | 1 | No | Yes |
| P05 | 8 | No | Yes |
| P06 | 0 | No | Yes |
| P07 | 14 | No | Yes |
| P08 | 1 | No | Yes |
| P09 | 12 | No | No |
| P10 | 0 | Yes | Yes |

**Table S3. Summary of behavioral performance for the MRI experiment.** Mean (SD) percent correct across runs was 97.75 (3.59) %. All but two participants had an accuracy above 99%. The two exceptions [P07: 88.08 (5.02) %; P10: 91.42 (11.43) %] may have been caused by a combination of the insensitive button box and slight sleepiness. Nevertheless, through careful inspection of their fMRI data, we found no obvious difference in either data quality or general tuning patterns between the two exceptions and the remaining participants. Their data were therefore kept in the fMRI analyses.

| Participant | Mean (%) | SD (%) |
|---|---|---|
| P01 | 100.00 | 0.00 |
| P02 | 99.83 | 0.56 |
| P03 | 99.33 | 1.52 |
| P04 | 99.83 | 0.56 |
| P05 | 99.92 | 0.41 |
| P06 | 100.00 | 0.00 |
| P07 | 88.08 | 5.02 |
| P08 | 99.92 | 0.41 |
| P09 | 99.17 | 1.31 |
| P10 | 91.42 | 11.43 |
| Group | 97.75 | 3.59 |
